# Circadian PERIOD proteins sculpt the mammalian alternative splicing landscape

**DOI:** 10.1101/2024.12.23.630108

**Authors:** L Chikhaoui, K Mamgain, M Seki, C Blanco, F Sassolas, E Folco, D Sery, Y Suzuki, B Ananthasubramaniam, K Padmanabhan

**Author notes:** Genomics Group, Institute of Food Safety & Analytical Sciences, Nestlé Research, Vers-chez-les-Blanc, Lausanne, Switzerland (L Chikhaoui). co-first author (L Chikhaoui, K Mamgain).

## Abstract

Mammalian circadian oscillators are driven by a transcription-translation feedback loop where CLOCK:BMAL1 activity is repressed by the PER:CRY complex. While transcriptional regulation by PER is well established, the role of circadian feedback in co- and post-transcriptional processes remains unclear. Here, we used Nanopore long-read direct RNA sequencing (dRNAseq) and quantitative mass spectrometry (qMS) to uncover a critical function of PERs in alternative splicing (AS) regulation in the liver. Our expanded long-read transcriptome revealed significant changes in rhythmic expression of annotated transcripts, novel isoforms of known genes, and previously unannotated genes, with widespread perturbations in Per1^-/-^;Per2^-/-^ (PerKO) livers. Rhythmic AS events were restricted to a distinct subset of transcripts, and splicing entropy - a metric of AS complexity - displayed oscillations in only a limited number of pathways, primarily those associated with glucose homeostasis and cellular responses to insulin. In PerKO livers, however, we detected increased isoform complexity and altered splicing entropy across a broad range of pathways linked to cell growth, morphogenesis, ER-associated degradation (ERAD), insulin response and histone methylation. Biochemical analyses and qMS data indicate that these changes are not due to mis-expression of splicing factors, but rather stem from altered nuclear abundance and chromatin retention of a few Serine-Arginine-rich splicing factors (SRSFs). In particular, SRSF3 acts proximal to the core-clock by defining both the period and amplitude of cellular rhythms. Our findings highlight a critical role for PER proteins in shaping the circadian liver proteome by integrating rhythmic transcription with the regulation of a complex and dynamic splicing landscape.

## Introduction

Circadian clocks are ubiquitous timekeepers that synchronize physiology, metabolism and behavior with the solar day. Cell-autonomous circadian oscillators drive daily rhythms in gene expression and protein synthesis to help build and maintain a homeostatic relationship with the external environment. Over two decades of genetic and proteomic screens have helped identify many components of the core clock and helped build a model of the regulatory network^1^. Within mammalian cells, a BMAL1:CLOCK transcription factor heterodimer drives the expression of the repressor complex components *Per* and *Cry*, which assemble in a ∼1MDa complex to repress BMAL1:CLOCK function on chromatin^2–5^. Because negative feedback inhibits new PER and CRY synthesis, PER and CRY protein concentrations decline via turnover, resulting in the termination of negative feedback and re-initiation of a new molecular cycle. This transcriptional-translational feedback loop operates in almost all cells and tissues to generate stable, self-sustaining molecular rhythms with an intrinsic period close to 24 hours. BMAL1:CLOCK via promoter E-box element recruitment drives the additional rhythmic transcription of a fraction of expressed genes in a tissue-specific manner. These transcripts serve as molecular outputs of the clock to control myriad cellular processes and typically do not participate in establishing the negative feedback loop. A second regulatory loop involving the orphan nuclear receptors Rev-erbα and RORα stabilizes the expression of Bmal1 via RORE elements^6^. The cis-acting elements of the core and accessory loops together promote precisely phased circadian transcription at clock output genes to establish the tissue proteome.

In addition to transcription, the regulation of co-transcriptional and post-transcriptional pathways including alternative splicing (AS) represents a critical layer of gene expression that shapes tissue proteomes and physiology. Most AS events occur co-transcriptionally, coordinated with elongating RNA Polymerase II (Pol II) on chromatin, and contributes to proteome diversity by generating multiple coding and non-coding isoforms from a single gene^7^. The splicing process is tightly regulated at nearly every step by Serine/arginine-rich splicing factors (SRSFs) - critical trans-acting factors that regulate both splice site recognition and spliceosome assembly^8^. Exon-bound SRSFs can act to enhance splicing while intron-binding often suppresses splicing. Beyond their co-transcriptional roles, SRSFs also influence mRNA translation as well as nonsense-mediated decay pathways to impact cellular protein composition. Their activities are further modulated by phosphorylation and dynamic shuttling between the nucleus and cytoplasm, adding complexity to the regulatory network governing mRNA fate^9^.

Circadian regulation of AS has been inferred from short-read sequencing data, revealing temporal control over splicing in mammalian tissues. For instance, time-of-day-dependent AS events have been observed in the pancreas^10^, while exon-array analyses in the liver identified “circadian” exons within key clock genes such as *Clock*, *Npas2*, and *Rev-erbα*. The latter finding suggested a bidirectional feedback loop in which the circadian clock regulates AS, and splicing decisions, in turn, influences the core clock^11^. A notable variation of this theme is the alternative splicing of *U2AF26,* which generates a variant that interacts with PER1 to modulate its stability^12^. Similarly, the RNA-binding proteins PSF and NONO, known components of the PER complex, are also known to regulate alternative splicing ^3,13,14^. However, the impact of PER-mediated circadian negative feedback on global AS landscapes remains unexplored.

Current mammalian circadian transcriptome datasets largely rely on short-read RNA sequencing (RNAseq) of PCR-amplified cDNA, which introduces biases during RNA extraction and amplification. Moreover, short-read analyses infer isoform expression probabilistically, limiting precision in transcript-level quantification^15^. To overcome these limitations, we employed Nanopore-based direct RNA sequencing (dRNASeq)^16^ on intact mRNA to generate isoform-level expression profiles in the murine liver over circadian time and in PerKO tissues and investigated the mechanism underlying AS changes using quantitative mass spectrometry. Our approach expands the transcriptome landscape in the murine liver and reveals isoform rhythms in newly identified transcripts from annotated genes as novel genes. Circadian regulation of AS spans all types of splicing events, which allowed us in turn to compute splicing entropy-first, to formalize the observed complexity of AS events and second, to pinpoint pathways specifically affected in PerKO livers. Quantitative mass spectrometry (qMS) revealed significant upregulation of a subset of Serine/Arginine-rich splicing factors (SRSFs) in the PerKO nucleoplasm. Biochemical fractionation of nuclei further demonstrated aberrant retention of Serine/Arginine-rich splicing factors (SRSFs) on PerKO chromatin, unlike wildtype livers, implicating co-transcriptional splicing mechanisms in these changes. These findings reveal that the circadian clock orchestrates pathway-specific mRNA oscillations through AS regulation. And that disruption of PER-mediated negative feedback changes the composition and localization of SRSF proteins, reshaping the AS network and the splicing landscape in the liver.

## Results

### Long-read direct RNAseq expands the murine liver transcriptome

To analyze the circadian transcriptome using dRNAseq, we sequenced Poly(A)+ enriched RNAs liver RNA from wild-type C57BL6 (4hr resolution, circadian times CT, CT0-CT20) and PerKO (CT8 and CT20) mice on the Oxford Nanopore PromethION platform (Supplemental Fig. S1A). We generated a total of 69 million high-quality long reads with an average read length of about 1000 nt that yielded 12855 expressed genes (Fig.1A and Supplemental Fig. S1A) covering >90% of the known expressed liver transcriptome^17^.

**Figure 1.**
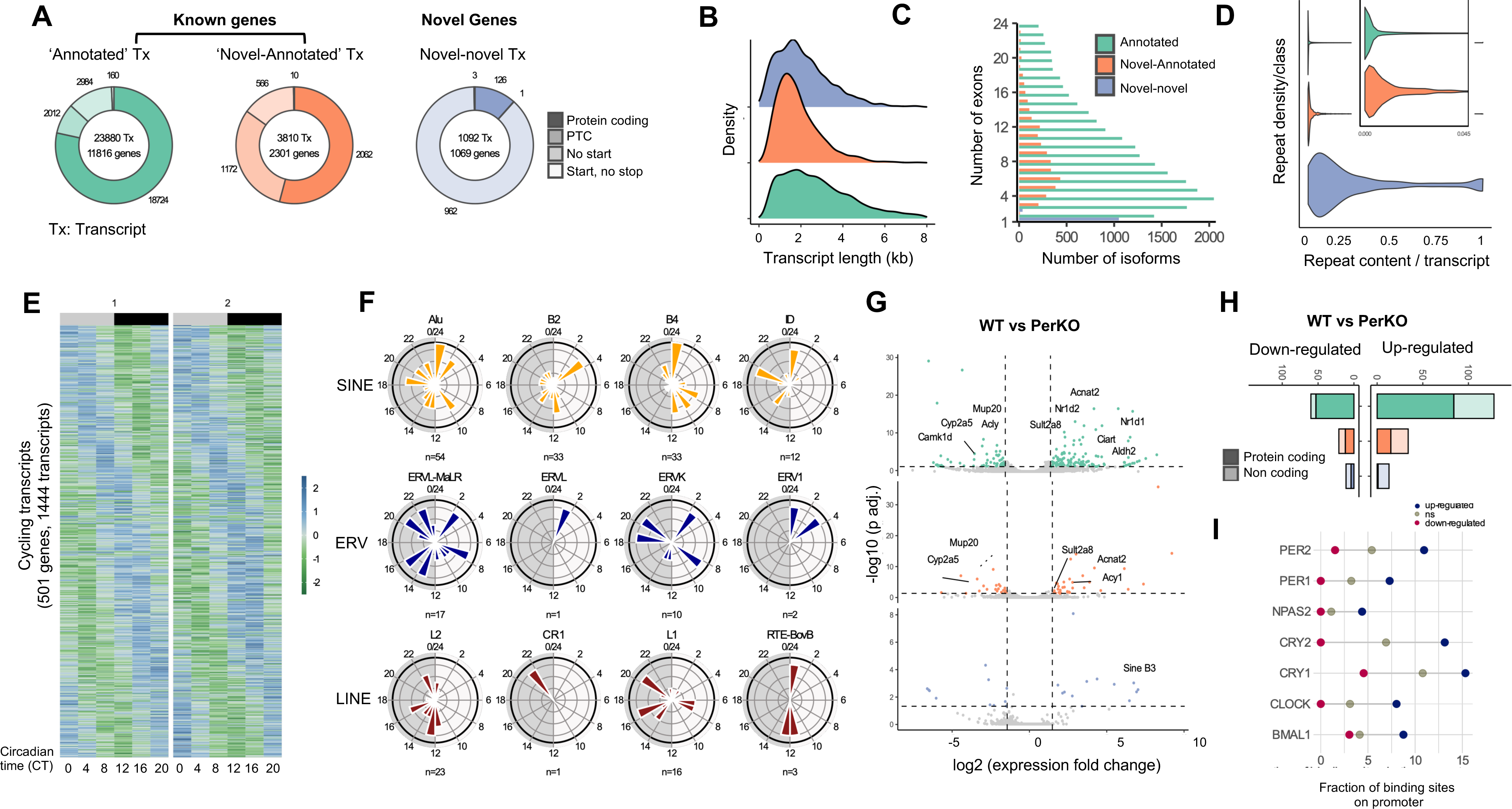
Direct RNAseq expands the known circadian and clock-disrupted liver transcriptome. **A)** De-novo transcriptome generated using FLAIR showing expressed transcripts (Tx) divided into three annotation classes. Annotated(green) and novel-annotated(orange) coming from known genes, and novel-novel(blue) coming from ‘novel’ genes. Donut plots show productivity, represented by shade for every transcript within a class. From darker to light shade for-productive, premature termination codon, no start codon, has start but no stop codon. Total number of transcript (Tx) and genes are shown in the centre, and numbers corresponding to the different productivity are displayed on the edge. **B)** Density plots showing transcript length (in kb), split and colored by annotation class. **C)** Number of exons per transcript, coloured by annotation class. **D)** Repeat content distribution per transcript in each annotated class. Zoomed in inset for the annotated and novel annotated class. **E)** GLM rhythmicity for circadian analysis. Heatmap showing expression of transcripts over circadian time (CT0-CT20). Each row represents a transcript, ordered by phase plotted are all expressed isoforms of genes that have >1 isoform expressed and at least 1 significantly cycling isoform. Color corresponds to the relative difference to the median intensity values. 1 and 2 represent two biological replicates. Colored bars on top represent subjective day (grey) or subjective night(black) **F)** Rhythmicity analysis of TE-containing transcripts (Yellow: short-interspersed nuclear elements SINE, Blue: endogenous retroviral class (ERV) and red: long-interspersed nuclear element LINE family) **G)** Fold change in transcript expression between wildtype and PerKO livers at CT20. Colored dots indicate significantly differentially expressed transcripts (multiple testing by stageR, p.adj < 0.05 and absolute log2FC > 1.5). Colored and split by annotated class. **H)** Number of transcripts up or down regulated in PerKO livers in comparison to WT livers. Each bar represents an annotated class and is shaded for productivity information **I)** Fraction of non-significant (grey), up (blue) or down (red) regulated transcripts with a binding site for core clock proteins in their promoter. Core-clock protein binding sites (Koike et al, 2012) targeting mis-expressed loci.

Long-read sequencing enables accurate estimation of transcript isoform abundance, rather than merely aggregate gene-level expression, particularly for genes with a high number of similar isoforms^18^. Using FLAIR^19^ with stringent parameters we reconstructed the de novo isoform-level transcriptome of polyadenylated RNA from mouse liver tissue. This approach generated a reliable set of both previously annotated and novel unannotated isoforms (Fig. 1A). Approximately 77% of the reads mapped to annotated isoforms derived from coding and non-coding genes. Novel isoforms of annotated genes (*novel-annotated* class) accounted for a further 18% of the total mapped reads. And the remaining (∼5%) reads mapped to genes that did not show any overlap with annotated RNA (*novel-novel* class) (Fig. 1A, Supplemental Fig. S1B). All three transcript classes exhibited comparable expression levels, (Supplementary Fig. S1C). While protein productivity information is available for the latest annotated transcriptome, FLAIR additionally allowed us predict protein productivity on our novel classes of transcripts based on the presence of an open reading frame, a start and/or a stop codon. The result (Fig. 1A) showed that while the majority of the known transcripts encode functional proteins, only about half of the novel-annotated transcripts are productive with most of them retaining a premature termination codon or lacking a canonical start codon. Furthermore, of the nearly 1069 novel genes that were discovered (Fig. 1A), the vast majority are mono-exonic (Fig. 1C) and only about 8% encode putative novel proteins. Thus, approximately 40% of novel transcript isoforms contribute to proteome diversity in mouse liver tissue, while the majority likely represent non-coding elements or may fulfill regulatory roles.

A singular advantage of dRNASeq is the enhanced ability to detect and annotate transcripts containing repetitive sequences derived from transposable elements (TEs), which are widely dispersed throughout the mammalian genome. The majority of novel-annotated and novel transcripts contains less than 5% of TEs, but the novel-novel transcripts include greater TE content (Fig. 1D). Indeed, newly discovered transcripts are enriched for repeats compared to the other classes and we defined 33 full-length (>80% of repeat sequence over 80% of the transcript) transcripts of TE origin, almost all of which were among the novel-novel (16) and annotated (14) transcripts. We further classified these full length TE transcripts according to Repbase-defined annotations^20^ (Supplementary Fig. S1D). We identified snRNA families in our novel-annotated transcripts, new families of LTR genes (RLTR4), IAP’s and LINEs in the novel-novel class (Supplementary Fig. S1D,E) with varying degrees of expression.

### Direct RNAseq expands the repertoire of liver transcripts under circadian control

We applied GLM-based cosinor analysis^21^ to assess 24-h oscillations in transcript rather than gene abundance across the newly annotated transcriptome. Unlike third-generation short-read sequencing, dRNASeq enabled rhythm analysis to cover all detected isoforms of cycling genes. Using stringent criteria for cyclicity, we identified 770 oscillating transcripts originating from 501 genes with a 24hr periodicity (Fig.1E). Visualization of all isoforms associated with these rhythmic genes (1444 transcripts) revealed that coding and non-coding isoforms exhibit oscillatory expression, albeit with low statistical significance in some cases (Fig1E). Circadian transcription in the liver predominantly peaked during transitions between subjective day and night, as well as between subjective night and day. Interestingly, non-coding transcripts exhibited higher abundance during the subjective night (Supplementary Fig. 1F, 1G). To explore whether full-length transposable element (TE) sequences displayed rhythmic expression, we analyzed their expression profiles. Both annotated and novel classes of LINE, SINE, LTR, and ERV families demonstrated circadian expression, with their peak expression phases plotted in Fig. 1F. Transcripts containing repeat sequences were predominantly expressed during transitional phases of the circadian cycle, with LINE families, in particular, peaking primarily during the subjective night (Fig. 1F).

Nuclear accumulation of PER and its role as a transcriptional repressor on chromatin peaks at CT20. To investigate the impact of PER loss at this critical time point, we performed a differential analysis of isoform expression in wildtype and PerKO livers. Loss of PER function resulted in the de-repression of core-clock targets (Fig. 1G-I). A total of 267 transcripts originating from 202 genes, encompassing annotated and newly annotated genes, were significantly differentially expressed in PerKO livers (p.adj < 0.05 and absolute log2FC > 1,5), with the majority being upregulated. Pathway enrichment analysis of these differentially expressed transcripts showed significant (p.adj < 0.05) enrichment of genes involved in pyruvate metabolism, regulation of IGF factor binding, uptake and transport, Chylomicron assembly and nuclear receptor transcription (Fig.1G, Supplemental Fig.S1H). Notably, most upregulated transcripts are targets of the core-clock transcription regulatory complex (Fig. 1I), including annotated transcripts for *Nr1d1* and *Nr1d2* and *Dbp* as well as novel isoforms for genes such as *Acnat2* and xenobiotic metabolism enzymes (*Cyp2a5 and Sult2a8*). A large fraction of the upregulated transcripts in the novel-annotated category included members of the major urinary protein (MUPs) and serine protease inhibitor (Serpin) families. MUPs play a role in stabilizing the release of pheromones in urine and regulate social and sexual behaviors. Of particular interest, Mup20 (also known as Darcin) is exclusively expressed in males, is inherently attractive to females and is associated with reinforcement of learning behaviors linked to mate selection and mating^22^. Among the ∼15 *Mup* genes that exhibited differential expression, *Mup20* was uniquely downregulated in PerKO livers. Finally, the novel-novel transcript category, showed upregulated RLTR4, SINEB3 and composite LTR/L1 familes transcripts in PerKO livers suggesting circadian regulation of their expression.

### Isoform-specific temporal dynamics in gene expression

Among the cycling transcripts, we observed that most core-clock and clock-output genes generate protein-coding isoforms that oscillate in phase (Fig. 2A, Supplementary Fig. S2A). Several loci exhibited three or more in-phase cycling isoforms including *Gpcpd1* (6), *Gsta3* (6) *Insig2* (7) *Pkbkf3* (5) *Mcm10* (4) *Gsta2* (4), *Dhrs3* (4), *St5* (5) *Hsd3b7* (6), *Tspan4* (5), *Hacl1* (9) and *Nudt7* (15). For *Bmal1*, *Per1* and *Ciart*, we identified three transcript isoforms that differed only in their 5’UTR sequences but otherwise encoded identical proteins. Among these, we identified a novel-annotated isoform for *Ciart,* a premature termination codon-containing ncRNA isoform for *Noct* and *Pfkfb3* which encode Nocturnin and Phosphofructokinase -key liver metabolic enzymes with circadian functions. Circadian transcription factors *Tef* expressed three protein coding isoforms while the *Dbp* locus additionally expressed a non-coding RNA. Transcripts originating from the Fatty acid synthase (*Fasn*) gene and its upstream regulatory TF, the Sterol Regulatory Element-binding Protein 1 exhibited 5 or more isoforms that oscillated in phase with a majority classified as ncRNAs due to premature termination. Notably, exon skipping events at the *Srebp1* locus resulted in 2 protein coding isoforms that lacked most of the N-terminal activation domains of the transcription factor as well as part of the lamin A interaction domain. Similarly, *Fasn* generated a cycling isoform that lacked an N-terminal exon, likely impacting its 3D folding and beta-ketoacyl synthase activity.

**Figure 2:**
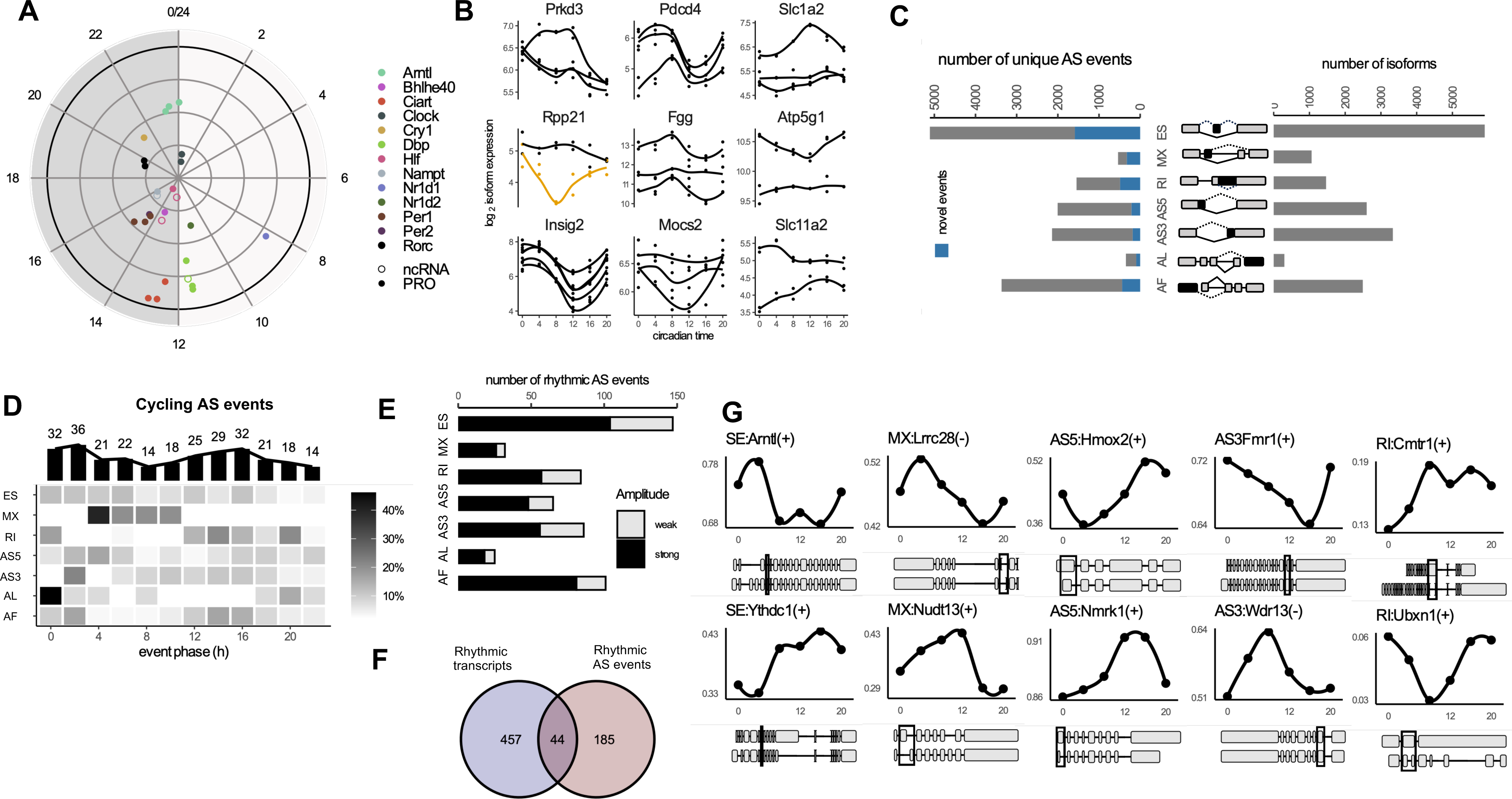
Transcript isoform dynamics in circadian livers. **A)** Phase distribution diagram representing 24 hours of a day. Each dot represents peak phase of all transcripts and their relative amplitude (from centre to periphery) for core-clock or key output genes. Filled circles represent productive isoforms and empty circles non-coding isoforms. **B)** Fold change (log2) in gene expression plotted over 24h for isoforms of genes that show differential transcript usage over circadian time. Showing productive (black) and non-productive (yellow) isoforms. **C)** Cartoon illustrates the alternative transcript generation event outputs of SUPPA analysis. Left panel shows the number of unique events present per event type, blue representing events that are detected because of presence of at least one novel isoform (novel events). Right panel shows number of isoforms involved in the respective event types. (ES, exon skipping; MX, mutually exclusive exons; RI, intron retention; A5S, alternate 5’ site; AS3, alternate 3’ splice site; AL, alternate last exon and AF, Alternate first exon) **D)** Heatmap for cycling AS events. Each row represents an event type, filled with the fraction of events peaking at the indicated CT phase. Bar plots on top show the total number of cycling AS event occurring at different time points. **E)** Bar plot showing number of cycling AS events classified as strong (black, amplitude >0.1) or weak (grey, amplitude <0.1) **F)** Intersection (middle) between genes that have rhythmic transcripts (left) and genes that have rhythmic AS events (right). **G)** Examples of cycling AS events [event type: gene name(strand)], showing cycling PSI values over circadian time. Boxes below show event coordinates depict the structure of the transcripts (one inclusive and one alternate isoform) for each example.

To investigate whether single gene loci can produce isoform-specific dynamics, we performed a differential transcript usage (DTU) analysis. This analysis identified circadian genes with isoform-specific dynamics, where some isoforms were arrhythmic or exhibited out-of-phase oscillations that produced arrhythmic isoforms or out-of-phase alternative cycling transcripts (Fig. 2B). For instance, *Prkd3, Slc1a2, Mocs2 and Atp5g1* produced one isoform with high-amplitude oscillations, while other isoforms maintained constant expression levels across the circadian cycle. The Rpp21 locus generated a high-amplitude rhythmic ncRNA isoform, while its coding isoform, which encodes a subunit of nuclear ribonuclease P, remained stably expressed. These results demonstrate that for a subset of expressed genes in the liver, the circadian clock can selectively regulate rhythmic transcript abundance for specific isoforms.

### Diverse co- and post-transcriptional events underlie liver isoform complexity

After reconstructing the liver transcriptome at the isoform-level, we used our annotation to identify and quantify individual alternative splicing (AS) events, revealing the complete AS landscape of our dataset (Supplementary Fig. S2B). We employed *Suppa*, a computational tool which enables detection and analyses of splicing events from multivariate biological samples, to identify splicing events at exons or introns that generate alternate transcripts from a given locus^23^. Our analysis classified these events into seven distinct categories: exon skipping (ES), mutually exclusive exons (MX), intron retention (IR), alternative 5’ splice-site (AS5) alternative 3’ splice-site (AS3) exons, alternative last exon (AL), and alternate first exon (AF). We also included alternate first exon (AF) usage that reflects Pol2 promoter choice driven isoform diversification. Of all the genes which display AS events, we focused our analysis on a subset where both the canonical and alternative isoforms were expressed (Supplementary Fig. S2B), ensuring that only events generating detectable alternative isoforms were considered. For example, an exon skipping event was included only if both isoforms (with and without the exon) were detected in our dataset.

ES was the most frequent event-type accounting for ∼5,000 events and producing alternative isoforms in over 50% of expressed genes (Fig. 2C). Alternate 5’ and 3’ alternate splice site choice followed, each affecting approximately one-quarter of expressed genes. AF, IR, and MX events contributed between 1,000 and 3,000 isoforms each, while AL events accounted for only a small fraction (3%) of transcriptome diversity. Since we detected many novel transcripts, we then estimated the number of novel AS isoform generation events. For most event types, the fraction of novel events ranged between 10% and 30% (blue bars). A notable exception was MX (mutually exclusive exons that never co-exist in the same transcript, a type of AS event rendered usually undetectable in short read sequencing), where more than half of the detected events were novel.

Next, to quantify the identified AS events, we calculated the percent-splice-in (PSI) value, which represents the fraction of a gene’s transcripts that include a specific AS event. Using this approach, we identified 281 AS events to be circadian in the WT murine liver characterized by rhythmic changes in PSI values (GLM analysis; Fig. 2D, examples plotted in Fig. 2G, Supplementary Fig. S2C). The majority of cycling AS events displayed high amplitudes with PSI changes exceeding 10% (Fig. 2E). When examining the temporal distribution of cycling AS events, exon skipping (ES) events were broadly distributed across the 24hr cycle. In contrast, AL events were notably absent during the early subjective night while IR events were predominantly observed during the night likely contributing to the higher abundance of non-coding transcripts during this period (Supplementary Fig. S1F). We then compared the rhythmically expressed transcripts and rhythmically spliced transcripts and found minimal overlap (Fig.2F). These findings suggest that distinct regulatory mechanisms are at play.

Since SINE and LINE transposable element (TE)-derived sequences have been shown to co-localize with alternative splice sites in the human transcriptome^24^, we investigated whether genetic loci with circadian AS events were associated with TE sequences. To test this, we compared TE distributions at these loci to a randomized control set, which then allowed us to evaluate if any of the observed AS events were linked to non-random TE distributions (Supplementary Fig. S3). Similar to findings in humans, SINE elements were found to be enriched across all cycling AS events, LTR elements specifically enriched at AF events while LINE elements were associated with ES and A5 events.

### Combinatorial AS events add to the isoform repertoire

Long-read sequencing, which spans entire transcripts, can unambiguously identify individual alternative splicing (AS) events and combinations of event types occurring within each transcript. Among the 9000 isoforms associated with only one AS event-type (Fig. 3A), the frequency of event types mirrored their overall prevalence, with ES being the most common (Fig. 2C). Approximately 10% of expressed transcripts displayed a combination of 2 or more AS events (Fig. 3A). Among isoforms that exhibited two different AS event-types, the majority involved ES in combination with RI, AF, AS5 and AS3. For combinations of 3 event-types, which were observed in 672 isoforms, these typically involved the same common event seen in simpler combinations (Fig.3A, inset). While ES events dominated the complex splicing landscape, transcripts associated with mutually exclusive exons (MX) in combination with other splicing events were primarily linked to novel isoforms identified in our analysis.

**Figure 3:**
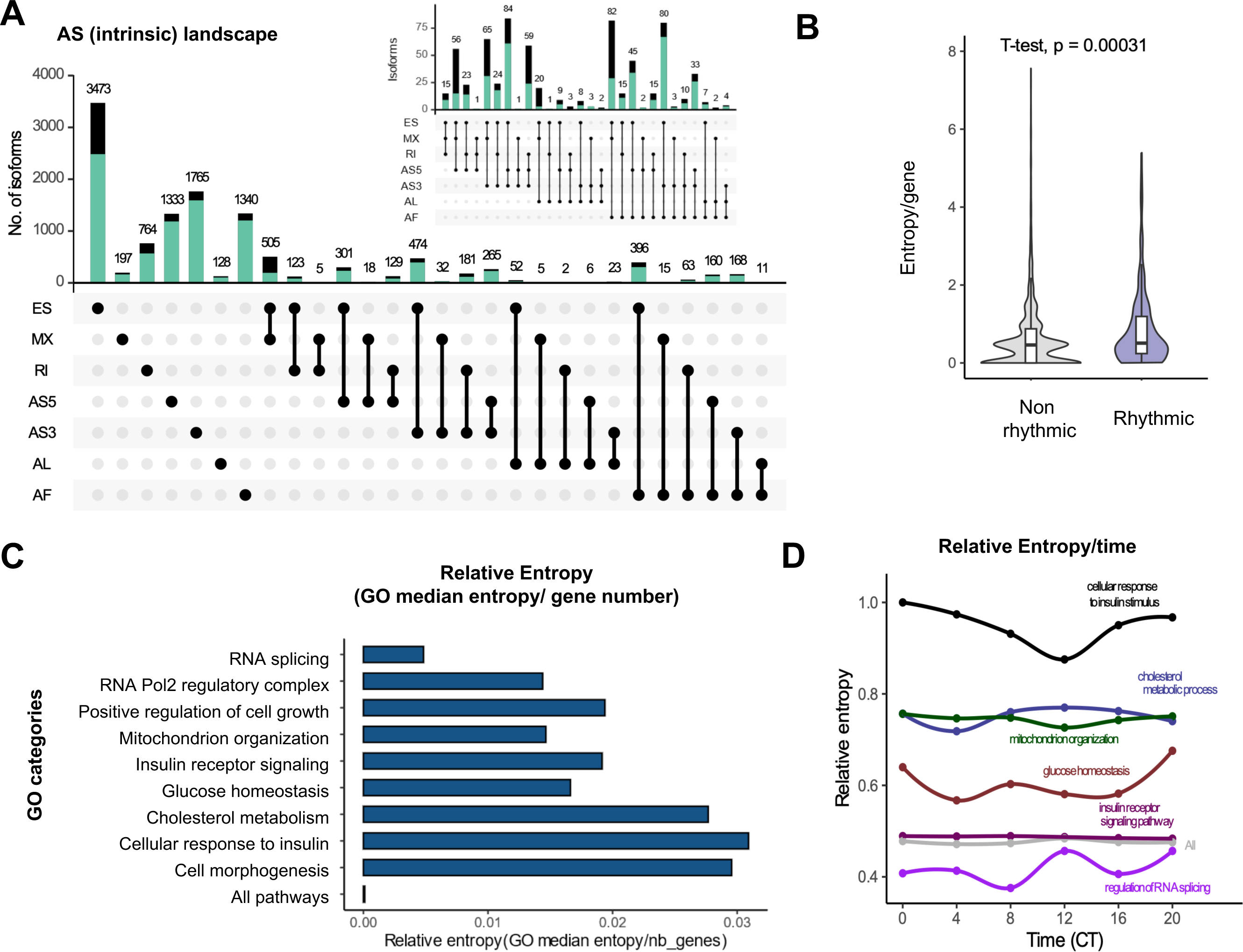
Complex AS events are widespread in the murine liver. **A)** Upset plot representing the total AS landscape of the expressed transcriptome. Total number transcripts with simple and complex splicing events (inset, >3 events in combination/transcript), green indicates fraction of annotated transcripts/category and black indicates novel-annotated isoforms **B)** AS entropy /gene for non-rhythmic and rhythmic gene GO sets **C)** Relative entropy (GO median entropy/maximum GO median entropy) for a subset of GO categories transcriptome wide combined for all WT samples **D)** Change in the relative entropy/time for select GO categories over time. ALL signifies 900 GO categories in total.

### Circadian dynamics in splicing entropy

Alternative splicing (AS) results in changes to the relative abundance of transcript isoforms produced by a single gene and it often coincides with complex patterns of splicing within the same transcript. To assess the contribution of all splicing outcomes to gene expression, we calculated splicing entropy, a PSI-dependent measure of AS complexity that evaluates the linkage relationship between alternative exons, independent of read depth^25^, for all time points in wildtype livers. Rhythmic genes exhibited significantly higher mean splicing entropy when compared to non-rhythmic sets (Fig.3B). To explore AS complexity in a broader biological context, we extended the entropy calculation from the gene level to the gene ontology (GO) level, analyzing splicing entropy across biological processes and cellular regulatory pathways (Fig.3C). Consistent with previous findings^26^, we found that fundamental cellular pathways and core biological functions showed low AS entropy (entropy < 0.5). These include pathways linked to cytoskeletal organization, nuclear pore assembly, DNA replication and transcription and dephosphorylation. In contrast, pathways linked to signaling cascades such as detection of phosphorylated amino acids showed high AS complexity (Fig.3C). Other pathways with elevated AS entropy in the liver included insulin signaling, peroxisome pathways and ATPase binding. We then investigated whether median splicing entropy within a GO pathway varied over a circadian day. Among the 9997 genes classified into 900 GO categories, only a small subset displayed time-dependent cycles in entropy. The GO cluster “Glucose homeostasis” and “Cellular response to insulin” as well as “Regulation of RNA splicing” exhibited clear circadian osciilations in splicing entropy (Fig.3D) while “Cholesterol homeostasis” and “Mitochondrion organization” pathways showed modest rhythms over 24 hours.

### Loss of PER disrupts alternative splicing

Since splicing entropy of key metabolic pathways are under circadian control, we wondered if the absence of a functional clock had limited impact on AS complexity. To address this, we analyzed changes in AS events between WT and PerKO livers at CT20 by quantifying differences in percent-splice-in (ΔPSI) values, with a threshold of adj. p < 0.05 and |ΔPSI| ≥ 10%. At the level of individual AS events, nearly all 6 AS events alongwith the differential use of AF positions were significantly altered in PerKO tissue (Fig.4A). Unlike gene expression that skewed upwards in PerKO livers, AS changes were bidirectional across all event categories (Supplementary Fig. S4A). Of the 971 events that we detected, skipped exons (ES) were the most prevalent category, followed by AF, A5 and A3 events. Interestingly, only a small fraction of differentially expressed genes were also differentially spliced (Fig. 4B). The weak correlation between differential gene expression and splicing changes in PerKO livers (Pearson’s R = 0.35, p < 2.2 × 10^-16^) indicates that distinct regulatory mechanisms are responsible for shaping the AS landscape in the absence of PER function (Fig. 4B, lower panel).

**Figure 4.**
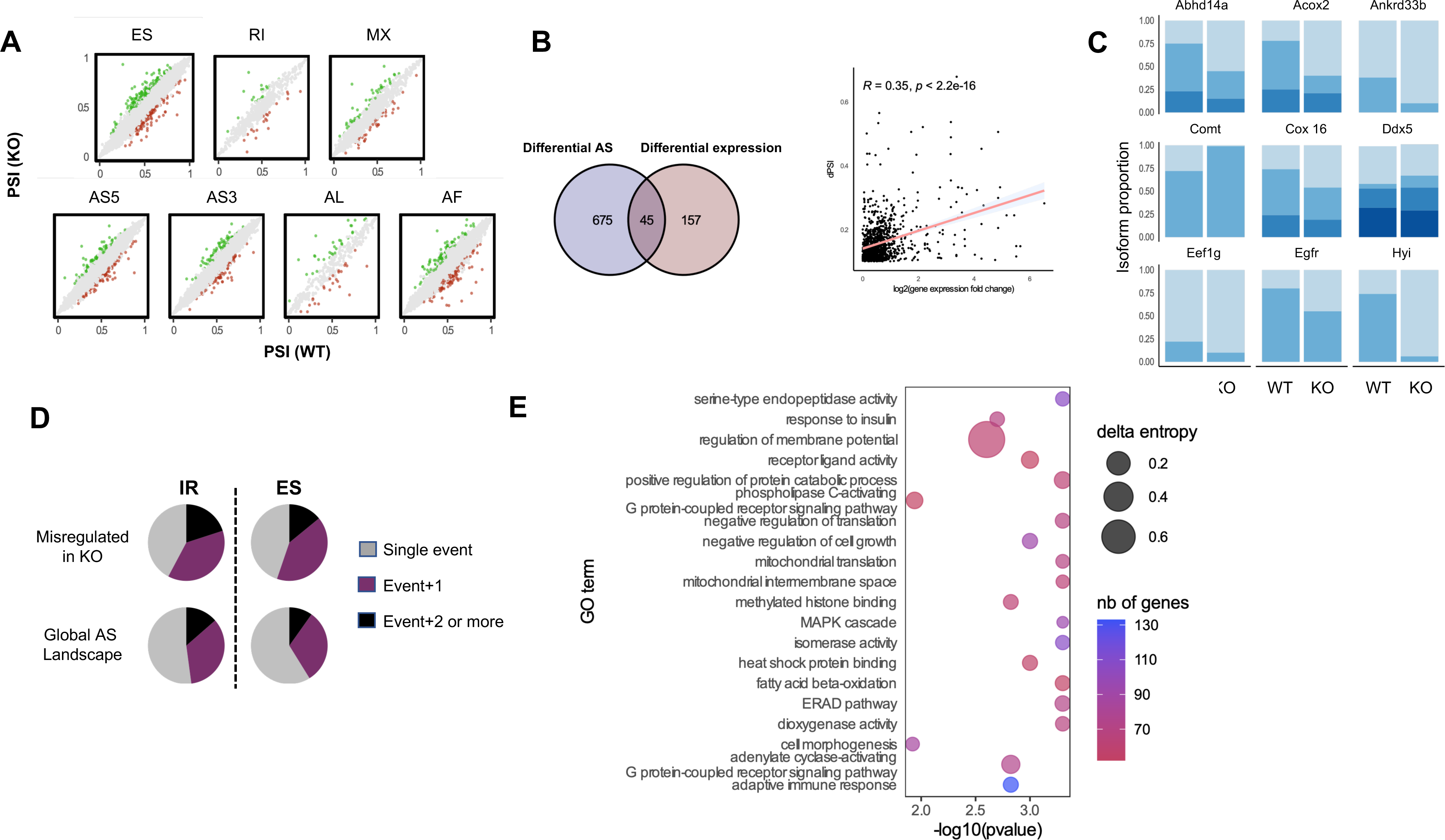
AS complexity in the absence of a functional clock in PerKO liver. **A)** Scatter plot showing PSI (percentage spliced in) values plotted for splicing events in WT and PerKO. Each dot represents a splicing event generated from SUPPA. Green and red dots correspond to significantly up-regulated and downregulated events in mutant (dPSI >/< 0.1 & pval <0.05) and grey indicates no change. **B)** Venn diagram showing intersection (middle) between genes that have differential splicing (left), and genes that have transcripts with differential expression (right) in PerKO*; lower panel* Pearson’s correlation analysis between log2FC and differential PSI (dPSI) between WT and PerKO **C)** Differential transcript usage at CT20 between wildtype and PerKO livers, each color within a bar indicates an individual transcript for a given gene. **D)** Splicing events per transcript in Global (all wildtype) transcripts and in transcripts with differential spicing (misregulated in PerKO, dpsi >0.1, pval <0.05). Fraction of all single unique AS events (exon skipping (ES) or intron retention (IR) are colored gray, fraction of transcripts that undergo either ES or IR plus an additional AS event is colored purple while those ES or IR containing transcripts with two or more additional AS events are black. **E)** Delta entropy (WT vs PerKO, CT20) analysis; plot shows selected GO categories (significant differential entropy, pvalue < 0.05, qvalue < 0.05). Sized by the delta entropy and coloured by number of genes in the GO category.

These findings suggest significant alterations in isoform usage in PerKO mice. Among the 156 isoforms from 76 genes exhibiting differential transcript usage in the PerKO liver, Fig. 4C highlights examples of notable isoform switches. Changes in isoform ratios varied widely, from drastic shifts as in the case of *Comt* (isoform 1: isoform 2 ratio changing from 0.3 to 0.01) to moderate changes as for *Egfr* and *Ankrd33b* (isoform1/isoform 2 ratio changing from 0.25 to 0.5 and 2.3 to 9 respectively).

Next, we investigated whether the misregulated transcripts at CT20 were enriched for specific AS events or combinations of events. Over 50% of all ES and IR splicing events in the global transcriptome were unique occurrences, while approximately 30% of IR and 25% of ES events were combined with at least one additional AS event (Fig. 4D, Global AS landscape). In stark contrast, transcripts that were significantly differentially spliced (WT vs PerKO, CT20) showed greater enrichment for complex splicing events (presence of more than one mis-regulated AS event within the same transcript) compared to the total expressed transcript pool (Fig.4D, Global AS landscape vs PerKO misregulated set). Transcripts containing ES or IR in combination with 2 or more additional events nearly doubled in PerKO tissues. Notably, complex splicing events (involving 3+ AS events), excluding MX events, were enriched in the Per KO misregulated subset (Supplemental Fig.S4B). These findings indicate that while simple AS events globally declined in PerKO livers, the reduced number of AS isoforms was accompanied by an increase in mis-regulated complex splicing events. Therefore, we examined splicing entropy at gene-ontology level and compared WT and PerKO transcriptomes. While the relative entropy remains unchanged for the near totality of GO categories, which encompasses ∼10000 genes, specific pathways including GPCR signaling, fatty acid beta-oxidation, cell growth and morphogenesis, methylated histone binding and insulin response showed significantly reduced entropy in PerKO tissues as expected (adj pval < 0.1; Fig.4E).

### PER loss redistributes SR proteins within the nuclear compartment

Transcripts that are up-regulated in PerKO livers represent direct targets repressed by the PER complex, while down-regulated loci likely reflect secondary effects resulting from the loss of circadian negative feedback (Fig.1F). To determine whether mis-regulated AS events in PerKO livers directly reflect gene mis-expression, we compared dPSI values of all splicing events with their respective transcript expression levels. Our analysis (Fig. 4B, *lower panel*) revealed only a weak correlation (Pearson’s coefficient, R = 0.35) indicating that changes in AS for most transcripts in Per KO livers were independent of changes in gene expression. This suggests that the absence of PER’s affects regulatory mechanisms that directly modulate alternative splicing pathways in the liver.

PER complexes are known to interact with RNA processing factors such as NONO/PSF and DHX9/SETX that are known to also play a role in co-transcriptional processes including splicing^3,4,13^. Given that small changes in the concentration of splicing regulatory factors can impact AS fidelity, we analyzed if PerKO phenotypes arose from mis-expression or mis-localization of splicing factors. Using quantitative mass spectrometry, we analyzed the nucleoplasmic and cytoplasmic fractions of liver tissue from wild-type and PerKO mice (CT8 and CT20 for WT; CT20 for PerKO) (Fig. 5A, schema in Supplementary Fig. S5A). Across these fractions, we detected approximately 4,000 proteins, of which 284 were differentially abundant (p adj. val < 0.1). In the cytoplasm at CT20, proteins involved in extracellular matrix regulation and lipid metabolism were up-regulated while fatty acid metabolism and corticosteroid response pathways were downregulated (Supplemental Fig. S5C). In PerKO, the cytoplasmic proteome reflected a clear enrichment in pathways linked to carboxylic acid and lipid metabolism (Supplemental Fig. S5D). Importantly, global changes in protein abundance, whether up- or downregulated, were largely independent of transcript expression levels across all analyzed pathways (Supplemental Fig. S5C,D lower panes).

**Figure 5:**
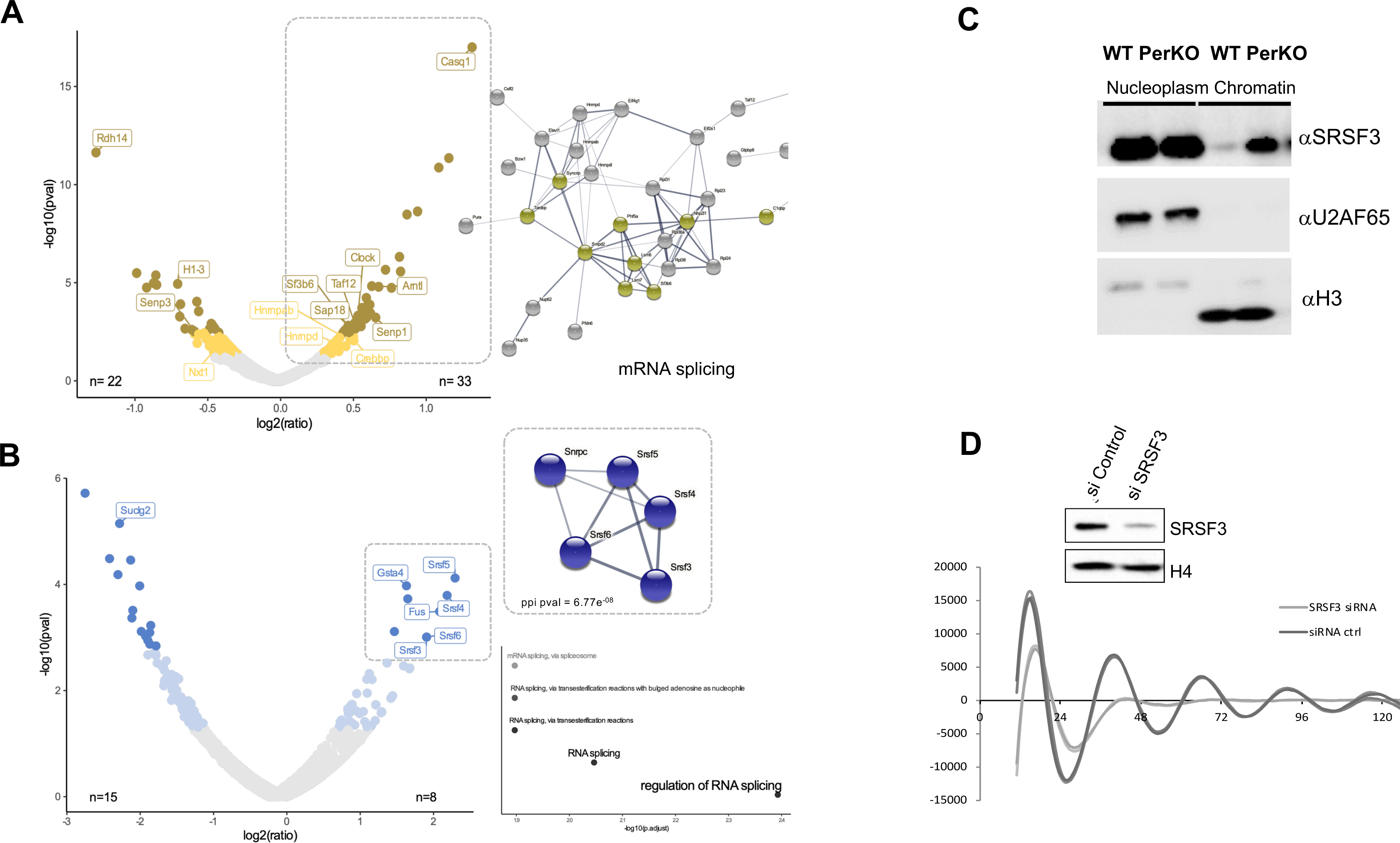
SRSF family proteins act in circadian negative feedback. **A)** Differential localization of proteins within WT nuclear compartments between CT8 and CT20. In yellow and ochre are proteins that significantly fractionate differentially (padj val < 0,1), *right panel* shows a string network analysis of the proteins, factors in yellow indicating proteins involved in mRNA splicing. **B)** Differential localization of proteins between WT and PerKO nuclear compartments between CT8 and CT20. In light and dark blue are proteins that show significant (padj val < 0,1) differential nuclear localization between the two conditions, *right panel* shows a string network analysis for proteins upregulated in the nuclear compartment in PerKO livers. **C)** Biochemical fractionation of nuclear protein into soluble nucleoplasm and chromatin-bound fractions. Protein extracts were probed for SRSF3, U2AF65 and histone H3 were used as loading controls for nucleoplasm and chromatin fractions respectively. **D)** Real time recording of luciferase signals over 5 days following knockdown of Srsf3 in U2OS fibroblasts harboring a Bmal1:luciferase reporter or a control scrambled siRNA. *Left panel* shows western blot for total SRSF3 protein in cell extracts; histone H4 is a loading control.

Among the 55 nuclear proteins enriched at CT8 in WT livers, we found the core-clock transcription factors BMAL1 and CLOCK as well as splicing factor SF3B6, regulatory protein Taf12, Hnrnpa/b and Hnrnpd and other proteins that play a role in mRNA splicing and metabolism (Fig.5A, Supplemental Fig. S6). In contrast, nuclear extracts from PerKO livers showed a significant accumulation of serine/arginine-rich splicing factors (SRSF) proteins (SRSF3, SRSF4, SRSF5, SRSF6), and FUS (Fig. 5B). Pathway analysis indicated that most upregulated proteins in PerKO livers were primarily involved in mRNA splicing, while down-regulated factors impacted metabolic processes (Supplemental Fig.S6A,B). The change in protein levels were independent of mRNA expression for most of these factors (Supplemental Fig.S6C). Thus, the aberrant AS landscape in PerKO liver is a complex phenomenon emerging from the accumulation of at least 6 SRSF proteins and FUS in the nuclear compartment coupled with change in chromatin architectural proteins (HMGB1 and HMGB2) that play additional roles in splicing and RNA metabolism^27^. We next examined the abundance and phospho-dynamics of proteins associated with the GO term “RNA splicing” in our dataset (Supplemental Fig.S7A). While mRNA expression levels of most of these factors didn’t change as a function of time (Supplemental Fig.S7A, left panel “expression”), their abundance or phosphorylation levels exhibited phase-specific peaks^28,29^ (Supplemental Fig.S7A, right panel “phase”). Thus, while the circadian regulation of AS is complex combinatorial event in wildtype animals, its disruption in the absence of circadian negative feedback likely results from a limited set of mis-localized SRSF proteins.

### SRSF3 acts as a core-clock component

Beyond a global regulation of AS programs in the liver, we then asked if any of the individual SRSF proteins that were enriched in the PerKO nucleus played a role in the core-circadian clock mechanism itself. Analysis of circadian ChIP-Seq datasets^30^ revealed that the SRSF3 locus was uniquely targeted by all core circadian regulators (Supplemental Fig.S7B). The AS function of SRSF3 is regulated by C-terminal phosphorylation domain of Pol2 and much like the PER complex, it has been implicated in regulation of Pol2 termination^4,31,32^. This function in turn requires its localization to nascent transcripts on chromatin. We therefore wondered if SRSF3 localization was altered within the nuclear compartment in PerKO livers. Liver nuclei from wildtype or PerKO livers were fractionated into nucleoplasm and chromatin fractions using urea-extraction buffer^33^. Nuclear fractionation of liver extracts revealed that while nucleoplasmic levels of SRSF3 remained unchanged, it was significantly enriched in the chromatin fraction in PerKO livers (Fig.5C, Supplemental Fig. S7C), indicating that SRSF3 is a clock-output gene that is clearly enriched on chromatin in the absence of a functional oscillator. To determine whether SRSF3 is required for cellular rhythmicity, we knocked down SRSF3 expression in U2OS cells expressing a Bmal1-Luc circadian reporter (Fig.5D, upper panel). Knockdown of SRSF3 resulted in damped rhythms followed by complete loss of circadian luciferase oscillations by day 3 (Fig.5D).

## Discussion

### dRNASeq analysis of a wildtype and a clock disrupted liver transcriptome

Over two decades of work has steadily expanded the circadian transcriptome beyond protein-coding genes to non-coding and regulatory RNAs in almost all mammalian tissues, in model and non-model organisms. Oscillatory transcription is linked to rhythms in histone modifications at promoters, gene bodies, and enhancers resulting in cycling mRNA, ncRNA and miRNA transcripts that play critical roles in the temporal compartmentalization of cellular and physiological processes. This is especially evident in the liver where the resident tissue clock regulates key hepatic functions such as lipid and carbohydrate metabolism, sterol biosynthesis and detoxification are regulated. Our work now reveals the extent to which both transcription and transcriptome diversity are clock-regulated phenomena and circadian disruption can re-distribute key alternative splicing factor hubs to dramatically impact the gene expression landscape.

Using Oxford Nanopore long-read dRNAseq, we generated isoform-level expression profiles in murine livers under circadian conditions in wildtype (WT) and clock-disrupted PerKO tissues. While minION platform data has been used in the past to annotate the liver transcriptome, the study suffered from low number (∼500,000) of reads. Previous studies using the MinION platform to annotate the liver transcriptome were limited by low read numbers (∼500,000). To overcome this, we sequenced each sample using a PromethION flow cell, ensuring sufficient read depth. Stringent cutoffs in FLAIR were applied to ensure the reliability of newly detected transcripts. Our AS analysis included only genes with both a primary transcript and an alternative isoform, effectively reducing the AS landscape by nearly half while increasing confidence in the detected isoforms and circadian dynamics. This approach expanded the transcriptome landscape in the wildtype (WT) liver and revealed isoform rhythms for core-clock genes, circadian output genes as well as expanded the circadian repertoire to TE-containing or full TE transcripts. In PerKO livers, long-read data revealed changes in AS events, including isoform switching in key metabolic genes and a previously underappreciated role for circadian regulation in mutually exclusive exons, intron retention, and exon skipping.

### SRSF proteins and clock function

The SRSF family, which includes 12 RNA binding proteins are trans-acting factors that regulate splice site recognition and spliceosome assembly during pre-mRNA splicing^36^. At steady state, they are localized in the nucleus and phosphorylation of the RS domain drives nucleo-cytoplasmic shuttling, the rates of which vary for different family members^37–39^. Changes in the abundance, localization, and activity of these proteins can affect splicing outcomes in rhythmic phenomena in plants^40,41^. In a similar vein, our qMS analysis revealed a significant upregulation of a subset of SRSF proteins specifically in the PerKO nucleus along with other RBPs such as FUS and TARDBP. Chromatin fractionation suggests that at least SRSF3 further accumulates on PerKO chromatin. Thus, the impact of SRSF mis-localization in PerKO tissues is likely to impact co-transcriptional pathways including polyadenylation and translation. Of all the SR proteins that are affected in PerKO livers, SRSF3 is a particularly strong candidate for a core-clock component. The SRSF3 promoter is targeted in a rhythmic manner by all core-clock proteins in the liver^30^, and we detected SRSF3 protein levels cycling in the tissue. In PerKO chromatin, we found a significant enrichment of SRSF3 that is likely to be regulating AS events co-transcriptionally. Moreover, real-time bioluminescence measurement of Bmal1-Luciferase expression following depletion of SRSF3 with specific siRNAs showed damped and eventual loss of circadian rhythms. The change in SRSF3 localization could be a simple downstream consequence of loss of PER-mediated chromatin organization^42^ or perhaps a more direct effect on active displacement of SRSF3 by the PER complex during negative feedback. Our experiments cannot distinguish between these two scenarios.

### Consequences of SRSF misregulation

Overexpression of SRSF proteins has been linked to higher risks to cancer progression and poor patient prognosis. High levels of SRSF3 has been reported in colorectal cancer, prostate and head and neck cancer. On the other hand, its depletion has predisposed mice to spontaneous HCC with aging. SRSF6 upregulation has been noted in colon and colorectal cancer, lung cancer and melanoma while SRSF5-7 were upregulated in small cell lung cancer. Upregulation of SRSF3 in particular has been shown to have a protective effect on HCC progression^43^ while its depletion promotes liver carcinogenesis^34^. Per KO animals are also predisposed to higher levels of HCC development^44^ and liver disease. In our dataset, we find SRSF6 and SRSF9 among cycling AS events, while expression of SRSFs doesn’t change significantly over circadian time or in PerKO livers. Thus, the greater abundance of 4 of these SR proteins and SRSF1 and SRSF7 (to a less significant extent) in the PerKO livers must result from regulation of their nucleo-cytoplasmic transport or translation of their mRNAs. PER proteins are thought to function as tumor suppressors and circadian clocks are often found to be disrupted in many cancers. Our work indicates an unexpected additional layer of regulatory mechanisms that may be operating in cancers where the disruption of clocks may drive the mis-localization of SRSF proteins, thereby impacting oncogenic progression. Transcriptome analysis of SRSF3 and SRSF1 liver-knockout mice have been described in the literature^34^,^35^. We re-analyzed Illumina short read transcriptome data from *Srsf1* and *Srsf3* KO livers by mapping it to our extended dRNAseq transcriptome annotation. All categories (annotated, novel-annotated and novel-novel) of transcripts could be detected in the public dataset. Furthermore, key circadian genes such as *Ciart, Usp2, Nr1d2* were significantly misregulated in *Srsf1* KO mice while *Rorc, Fus and Fasn* were mis-expressed in *Srsf3* KO livers (Supplemental Fig. S8) suggesting these knock-out tissues likely had disrupted rhythms.

Beyond its role in recruiting repressor complexes at core-clock gene and establishing chromatin states^1,6,45,46^, this study reveals that circadian negative feedback by PER complexes has a remarkable impact on the co- and post-transcriptional events in the nucleus. While the clock is necessary to generate transcript isoform diversity in a tissue, circadian disruption affects the localization and distribution of critical AS factors within the nuclear space and their targeting to chromatin. How dynamic epigenetic marks and alternative splicing pathways are coordinated in healthy tissues and otherwise disrupted in the absence of a functional clock remains to be explored.

## Acknowledgments

The work was supported by an ATIP Avenir installation grant, Projét Fondation ARC (PJA 20191209722), Pack Ambition Research AURA (2000671901) and an EU H2020 to KP. LC was funded by the Fondation Recherche Medicale (FRM) PhD fellowship. BA acknowledges grant AN1553/2-1 (Project No. 444137814) from the Deutsche Forschungsgemeinschaft (DFG). YS acknowledges JSPS KAKENHI Grant Number JP22H04925 (PAGS).

We would like to thank Marie-Paule Felder Schmittbuhl and Urs Albrecht for *Per1-/-; Per2 Brdm-/-* mice. We acknowledge the contribution of SFR Biosciences (UMS3444/CNRS, US8/Inserm, ENS de Lyon, UCBL) facility : PSF (Adeline Page and Frederic Delolme) for their help with mass spectrometry experiments. We would also like to thank Joel Richter, Virginie Faure and other members of IGFL for valuable inputs on the manuscript.

## Author Contributions

KP conceived and directed the project with LC, KM, YS and BA. LC, MS and KP conducted the experiments. LC and KM performed all the bioinformatics analyses with help from FS (TE analysis) and BA. Biochemical analysis, animal husbandry was performed by EF, CB and DS. KP wrote the manuscript with input from all authors.

## Materials and Methods

### Animals

6-week-old C57BL/6J male mice purchased from Charles River were bred with unlimited access to food (AL: ad libitum) and water in 12h:12h light-dark cycles conditions, for at least 2 weeks. One day prior to the killing, animals were transferred to constant darkness (DD) conditions. Liver samples were collected every 4 hours over 24 hours (Circadian Time CT) and snap froze into liquid nitrogen, stored at -80 °C until further use. *Per1^-/-^;Per2^brdm^* mice (Zheng *et al.* 2001) were subject to the same conditions and liver samples collected only at CT8 and CT20. Transcriptomic sequencing was performed on biological duplicates for each timepoint and proteomic analysis was performed on biological triplicates.

### Long-read sequencing and Data processing

#### Library preparation and Nanopore direct RNA sequencing

RNA from the two livers per time-point was extracted using standard Trizol RNA extraction. Poly(A) RNAs were isolated using μMACS mRNA Isolation kit (130-075-101). Five hundred nanograms was used for library preparation for direct RNA sequencing with SQK-RNA002 kit (Oxford Nanopore). Sequencing was performed using PromethION with one FLO_PRO002 flow cell per sample (Oxford Nanopore). Base-calling of the Fast5 data outputs from PromethION was performed using Guppy (v4.0.11) to convert into Fastq files.

#### Novel transcript discovery

Flair (v1.5) was used for novel transcripts discovery. Each sample reads were aligned onto mm39 reference mouse genome using native-RNA specific alignment parameters for minimap2 (v2.17) and samtools (v1.9). Flair collapse step default parameters were changed into more stringent filtering: we only retained novel transcripts with at least 20 supporting reads (--support 20) covering a minimum of 80% of the novel transcript (--stringent), and that are not a subset of an already annotated transcript (--filter nosubset) and map quality (mapq>10). Previously non-annotated transcripts passing these filters were added to the Ensembl annotate transcriptome to constitute our transcriptome of reference. Productivity of transcripts was predicted using Flair script “predictProductivity.py” based on the longest ORF for each transcript.

#### Quantification and differential transcript expression

The pipeline was established according to the most suitable analysis workflow described in Soneson *et al.* 2019. Reads were mapped onto the transcriptome of reference (containing novel transcripts) using Minimap2 (v2.17) with the nanopore specific preset “map-ont”, and only primary alignments were kept. Sorting and indexing of reads were performed using Samtools (v1.9). Isoform quantification was performed using Salmon (v1.5.2) in alignment-based mode and transcripts with an abundance of less than 20 reads across all conditions were filtered out. The number of reads per transcript was normalized to the total reads count of their corresponding sample and the batch effect was removed using limma.

#### Repeat identification

The mice genome repeats annotation was retrieved using UCSC Table Browser and intersected with our transcriptome of reference using bedtools (v2.30.0). Fraction of repeat content was calculated by the portion of repeats identified in each transcript. Transcripts containing more than 80% of a transposable element were considered as full-TE.

#### Circadian rhythmicity analysis

Multiple testing was performed using the stageR package in R and adjusted p-values <0.05 were considered significant. Alternatively, JTK cycle was used to assess the rhythmicity of transcripts, considered to be circadian when pvalue < 0.05. Differential isoform expression between WT and *Per1^-/-^;Per2^brdm^* mice at CT20 was performed using DESeq2.

#### Splicing analysis

Differential transcript usage was performed using StageR with false discovery rate set at 0.1 and p-values were adjusted to genes. Splicing analysis at the local splicing event was performed using SUPPA (v2.3): alternative splicing events were identified from the reference transcriptome containing novel isoforms and events quantification for each sample was performed using the batch corrected quantification for the transcripts considered to be expressed as described above. PSI values were used for circadian analysis using JTK cycle in WT. “diffSplice” module of SUPPA was used to identify differential splicing events between WT and *Per1^-/-^;Per2^brdm^* mice at CT20, the significance was calculated with empirical method and p-values were corrected by gene. Events with an adjusted p-value <0.05 and dPSI greater than 10% were considered significant.

Gene Ontology enrichment of differential alternative splicing events was calculated by comparing genes generating these events to genes subject to the same type of event as a background (for example all genes undergoing intron retention).

#### Entropy calculation

PSI values were averaged for each event across different replica and filtered to remove any mean psi values equal to 0. Gene entropy was then calculated according to the following formula: = - ∑_i_Ψ_i_log_2_Ψ_i_ where i is the PSI value of an event for a given gene.

### Proteomic analysis

#### Nuclei and cytoplasm fractionation

Each liver (in biological triplicate) was individually thawed on ice and then cut into pieces before homogenization in 3mL of PBS (supplemented with 0.5mM spermidine, 0.15mM spermine, 0.5mM DTT) with a dounce homogenizer. The homogenate was filtered on gauze mixed with 5mL of homogenization buffer (2.2M sucrose, 10mM Hepes pH 7.6, 15mM KCl, 2mM EDTA supplemented with 0.5mM spermidine, 0.15mM spermine, 0.5mM DTT) and re-homogenized. 20mL of homogenization solution was added to the mix and then layered on top of 9mL of cold cushion sucrose solution (2.05M sucrose, 10mM Hepes pH 7.6, 10% glycerol, 15mM KCl, 2mM EDTA supplemented with 0.5mM spermidine, 0.15mM spermine, 0.5mM DTT). Nuclei were pelleted by ultracentrifugation for 45min at 4°C at 25 000 rpm using a Beckman SW32 rotor. Supernatant corresponding to cytoplasmic fraction was collected while nuclei pellets were washed twice in NLB (10mM Hepes pH 7.6, 100mM KCl, 0.1mM EDTA pH 8.0, 10% glycerol supplemented with 0.5mM spermidine, 0.15mM spermine, 0.5mM DTT).

#### Protein extraction and preparation

Proteins from each fraction were extracted using 10% SDS – 3.5M β-mercaptoéthanol, quantified and 100ug of proteins per sample were used for mass spectrometry sample preparation. Proteins were digested using LysC/Trypsin (1/20) for 18 hours at 37°C, desalted on spin column C18 (Promega) and labeled with TMT 0.2 mg. Chromatin extracts were prepared according to Wuarin and Schibler, 1994.

#### Quantitative Mass spectrometry analysis

The samples were analyzed on a high-resolution orbitrap mass spectrometer in TOP15 HCD mode, R(MS2) : 45K. The samples were analyzed in technical triplicate. Data were reprocessed with PD2.5 software with the Sequest HT search engine against the Sprot Mus musculus database, and the addition of a contaminant bank, filtered to a 1% false positive rate. The quantification data from the 3 replicates were averaged and the p-value was calculated. Adjusted p-value <0.1 were considered as significant. Gene ontology enrichment analysis was performed on the whole mouse genome on ^30,31^. Network interaction visualization was created on https://string-db.org.

## References

1. Cox, K. H. & Takahashi, J. S. Circadian clock genes and the transcriptional architecture of the clock mechanism. J Mol Endocrinol 63, R93–R102 (2019).

2. Brown, S. A. et al. PERIOD1-associated proteins modulate the negative limb of the mammalian circadian oscillator. Science 308, 693–696 (2005).

3. Duong, H. A., Robles, M. S., Knutti, D. & Weitz, C. J. A molecular mechanism for circadian clock negative feedback. Science 332, 1436–1439 (2011).

4. Padmanabhan, K., Robles, M. S., Westerling, T. & Weitz, C. J. Feedback regulation of transcriptional termination by the mammalian circadian clock PERIOD complex. Science 337, 599– 602 (2012).

5. Ju, D. et al. Chemical perturbations reveal that RUVBL2 regulates the circadian phase in mammals. Sci Transl Med 12, eaba0769 (2020).

6. Papazyan, R., Zhang, Y. & Lazar, M. A. Genetic and epigenomic mechanisms of mammalian circadian transcription. Nat Struct Mol Biol 23, 1045–1052 (2016).

7. Carrocci, T. J. & Neugebauer, K. M. Emerging and re-emerging themes in co-transcriptional pre-mRNA splicing. Mol Cell 84, 3656–3666 (2024).

8. Braunschweig, U., Gueroussov, S., Plocik, A. M., Graveley, B. R. & Blencowe, B. J. Dynamic integration of splicing within gene regulatory pathways. Cell 152, 1252–1269 (2013).

9. Kretova, M., Selicky, T., Cipakova, I. & Cipak, L. Regulation of Pre-mRNA Splicing: Indispensable Role of Post-Translational Modifications of Splicing Factors. Life (Basel) 13, 604 (2023).

10. Marcheva, B. et al. A role for alternative splicing in circadian control of exocytosis and glucose homeostasis. Genes Dev 34, 1089–1105 (2020).

11. McGlincy, N. J. et al. Regulation of alternative splicing by the circadian clock and food related cues. Genome Biol 13, R54 (2012).

12. Preußner, M. et al. Rhythmic U2af26 alternative splicing controls PERIOD1 stability and the circadian clock in mice. Mol Cell 54, 651–662 (2014).

13. Kowalska, E. et al. NONO couples the circadian clock to the cell cycle. Proc. Natl. Acad. Sci. U.S.A. 110, 1592–1599 (2013).

14. Heyd, F. & Lynch, K. W. Phosphorylation-dependent regulation of PSF by GSK3 controls CD45 alternative splicing. Mol Cell 40, 126–137 (2010).

15. Jin, H., Wan, Y.-W. & Liu, Z. Comprehensive evaluation of RNA-seq quantification methods for linearity. BMC Bioinformatics 18, 117 (2017).

16. Garalde, D. R. et al. Highly parallel direct RNA sequencing on an array of nanopores. Nat Methods 15, 201–206 (2018).

17. Yu, Y. et al. A comparative analysis of liver transcriptome suggests divergent liver function among human, mouse and rat. Genomics 96, 281–289 (2010).

18. Pardo-Palacios, F. J. et al. Systematic assessment of long-read RNA-seq methods for transcript identification and quantification. Nat Methods 21, 1349–1363 (2024).

19. Tang, A. D. et al. Full-length transcript characterization of SF3B1 mutation in chronic lymphocytic leukemia reveals downregulation of retained introns. Nat Commun 11, 1438 (2020).

20. Jurka, J. Repbase update: a database and an electronic journal of repetitive elements. Trends Genet 16, 418–420 (2000).

21. Love, M. I., Huber, W. & Anders, S. Moderated estimation of fold change and dispersion for RNA-seq data with DESeq2. Genome Biol 15, 550 (2014).

22. Demir, E. et al. The pheromone darcin drives a circuit for innate and reinforced behaviours. Nature 578, 137–141 (2020).

23. Alamancos, G. P., Pagès, A., Trincado, J. L., Bellora, N. & Eyras, E. Leveraging transcript quantification for fast computation of alternative splicing profiles. RNA 21, 1521–1531 (2015).

24. Clayton, E. A. et al. An atlas of transposable element-derived alternative splicing in cancer. Philos Trans R Soc Lond B Biol Sci 375, 20190342 (2020).

25. Sterne-Weiler, T., Weatheritt, R. J., Best, A. J., Ha, K. C. H. & Blencowe, B. J. Efficient and Accurate Quantitative Profiling of Alternative Splicing Patterns of Any Complexity on a Laptop. Mol Cell 72, 187–200.e6 (2018).

26. Zhao, F., Yan, Y., Wang, Y., Liu, Y. & Yang, R. Splicing complexity as a pivotal feature of alternative exons in mammalian species. BMC Genomics 24, 198 (2023).

27. Sofiadis, K. et al. HMGB1 coordinates SASP-related chromatin folding and RNA homeostasis on the path to senescence. Mol Syst Biol 17, e9760 (2021).

28. Wang, J. et al. Nuclear Proteomics Uncovers Diurnal Regulatory Landscapes in Mouse Liver. Cell Metab 25, 102–117 (2017).

29. Robles, M. S., Humphrey, S. J. & Mann, M. Phosphorylation Is a Central Mechanism for Circadian Control of Metabolism and Physiology. Cell Metab 25, 118–127 (2017).

30. Koike, N. et al. Transcriptional architecture and chromatin landscape of the core circadian clock in mammals. Science 338, 349–354 (2012).

31. Cui, M. et al. Genes involved in pre-mRNA 3’-end formation and transcription termination revealed by a lin-15 operon Muv suppressor screen. Proc Natl Acad Sci U S A 105, 16665–16670 (2008).

32. Le Bozec, B. et al. Circadian PERIOD proteins regulate TC-DSB repair through anchoring to the nuclear envelope. Preprint at 10.1101/2023.05.11.540338 (2023).

33. Wuarin, J. & Schibler, U. Physical isolation of nascent RNA chains transcribed by RNA polymerase II: evidence for cotranscriptional splicing. Mol. Cell. Biol. 14, 7219–7225 (1994).

34. Sen, S., Jumaa, H. & Webster, N. J. G. Splicing factor SRSF3 is crucial for hepatocyte differentiation and metabolic function. Nat Commun 4, 1336 (2013).

35. Arif, W. et al. Splicing factor SRSF1 deficiency in the liver triggers NASH-like pathology and cell death. Nat Commun 14, 551 (2023).

36. More, D. A. & Kumar, A. SRSF3: Newly discovered functions and roles in human health and diseases. Eur J Cell Biol 99, 151099 (2020).

37. Cáceres, J. F., Screaton, G. R. & Krainer, A. R. A specific subset of SR proteins shuttles continuously between the nucleus and the cytoplasm. Genes Dev 12, 55–66 (1998).

38. Sapra, A. K. et al. SR protein family members display diverse activities in the formation of nascent and mature mRNPs in vivo. Mol Cell 34, 179–190 (2009).

39. Haward, F. et al. Nucleo-cytoplasmic shuttling of splicing factor SRSF1 is required for development and cilia function. Elife 10, e65104 (2021).

40. Staiger, D. & Brown, J. W. S. Alternative splicing at the intersection of biological timing, development, and stress responses. Plant Cell 25, 3640–3656 (2013).

41. Romanowski, A., Schlaen, R. G., Perez-Santangelo, S., Mancini, E. & Yanovsky, M. J. Global transcriptome analysis reveals circadian control of splicing events in Arabidopsis thaliana. Plant J 103, 889–902 (2020).

42. Tartour, K. et al. Mammalian PERIOD2 regulates H2A.Z incorporation in chromatin to orchestrate circadian negative feedback. Nat Struct Mol Biol 29, 549–562 (2022).

43. Wang, H. et al. Alteration of splicing factors’ expression during liver disease progression: impact on hepatocellular carcinoma outcome. Hepatol Int 13, 454–467 (2019).

44. Kettner, N. M. et al. Circadian Homeostasis of Liver Metabolism Suppresses Hepatocarcinogenesis. Cancer Cell 30, 909–924 (2016).

45. Zhu, Q. & Belden, W. J. Molecular Regulation of Circadian Chromatin. J Mol Biol 432, 3466– 3482 (2020).

46. Tartour, K. & Padmanabhan, K. The Clock Takes Shape-24 h Dynamics in Genome Topology. Front Cell Dev Biol 9, 799971 (2021).

