## Supplementary Figures and Legends for "Circadian PERIOD proteins sculpt the mammalian alternative splicing landscape"

### Supplemental Figures

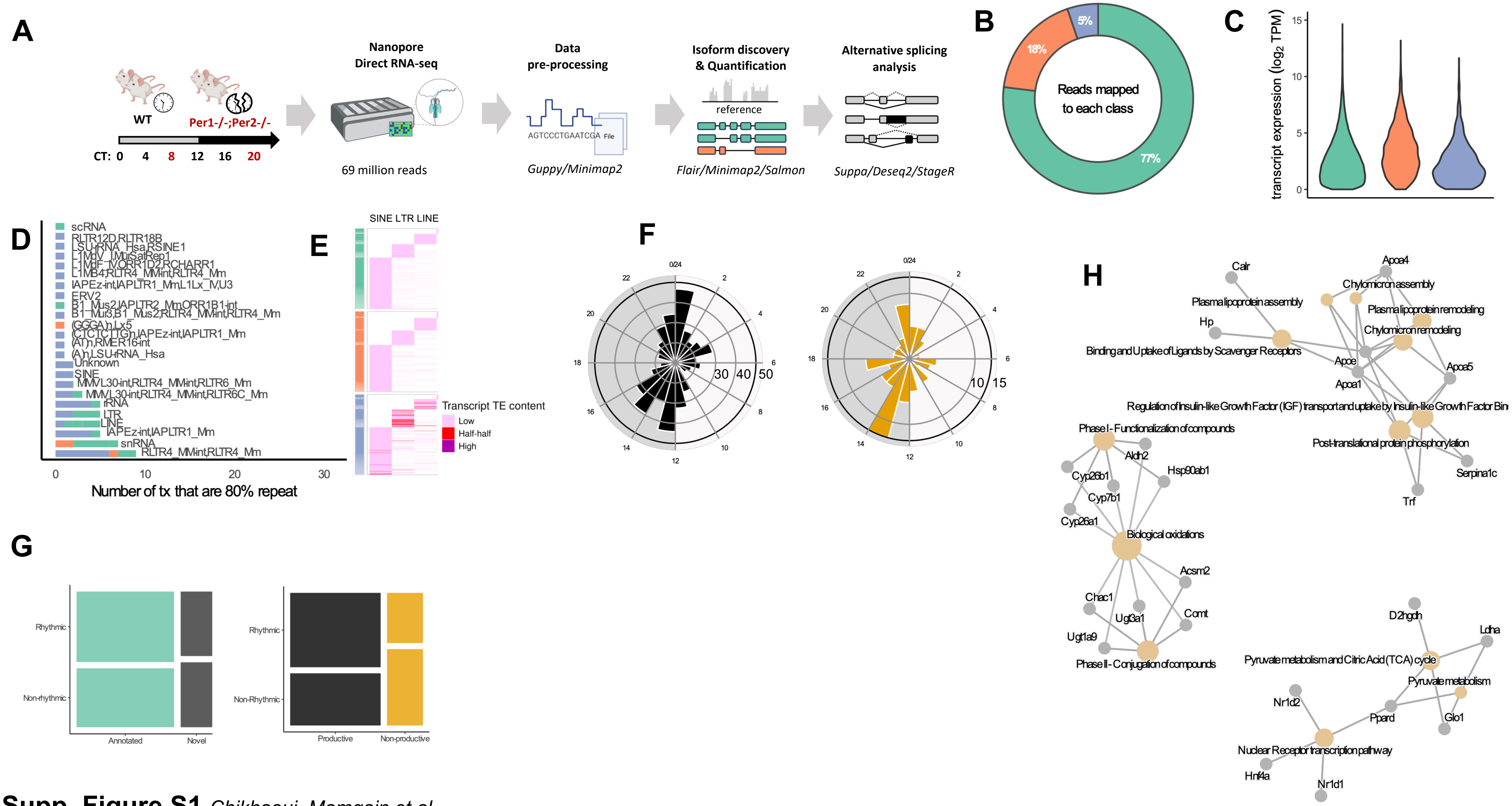


**Supp. Figure S1** *Chikhaoui, Mamgain et al*

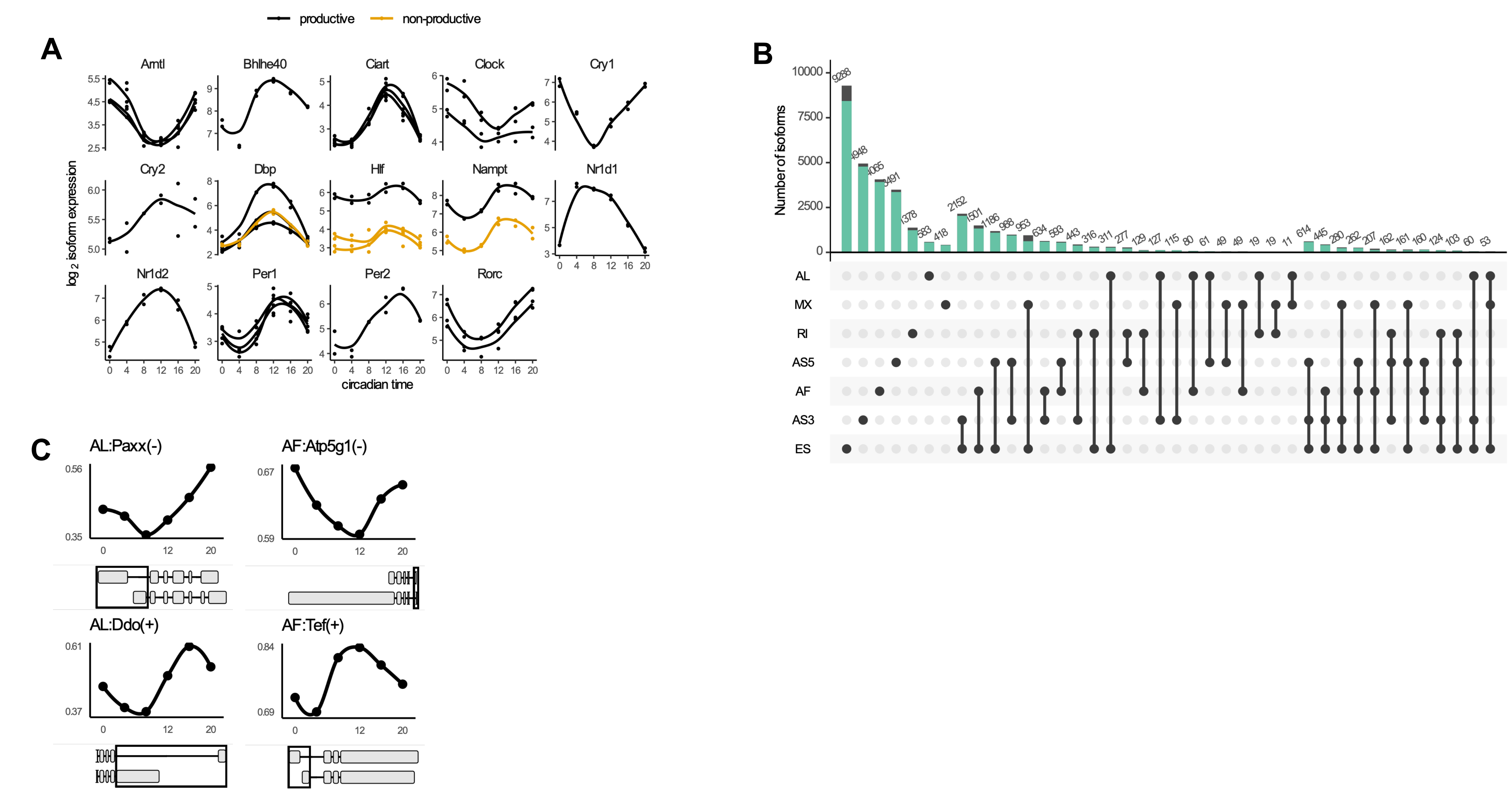


**Supp. Figure S2**

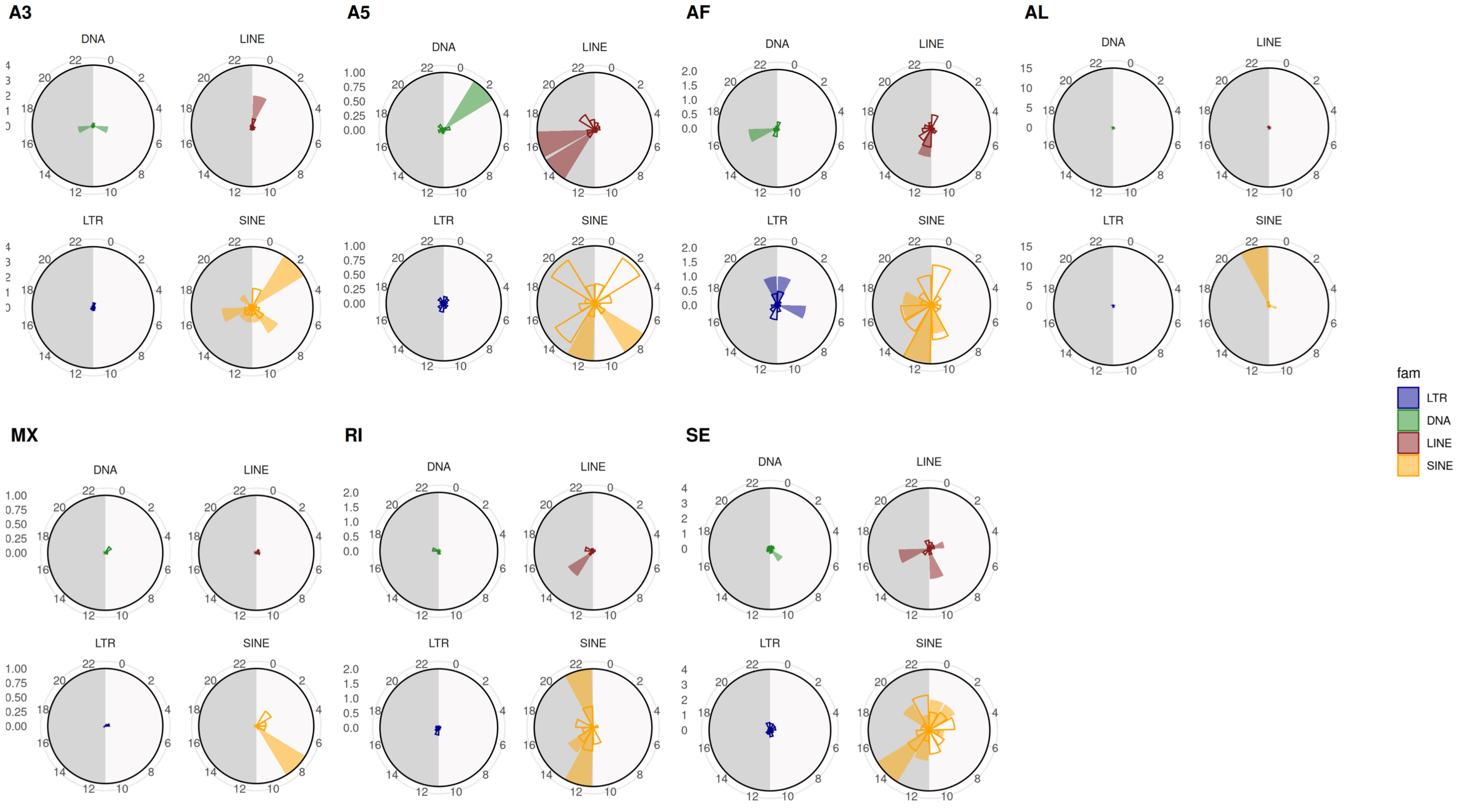


**Supp. Figure S3** *Chikhaoui, Mamgain et al*

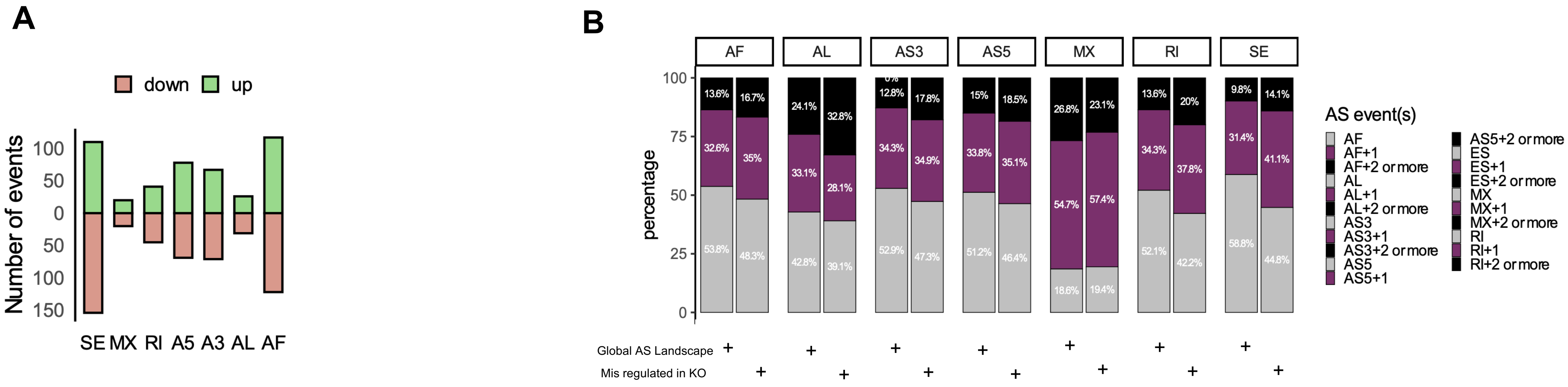


**Supp. Figure S4** *Chikhaoui, Mamgain et al*

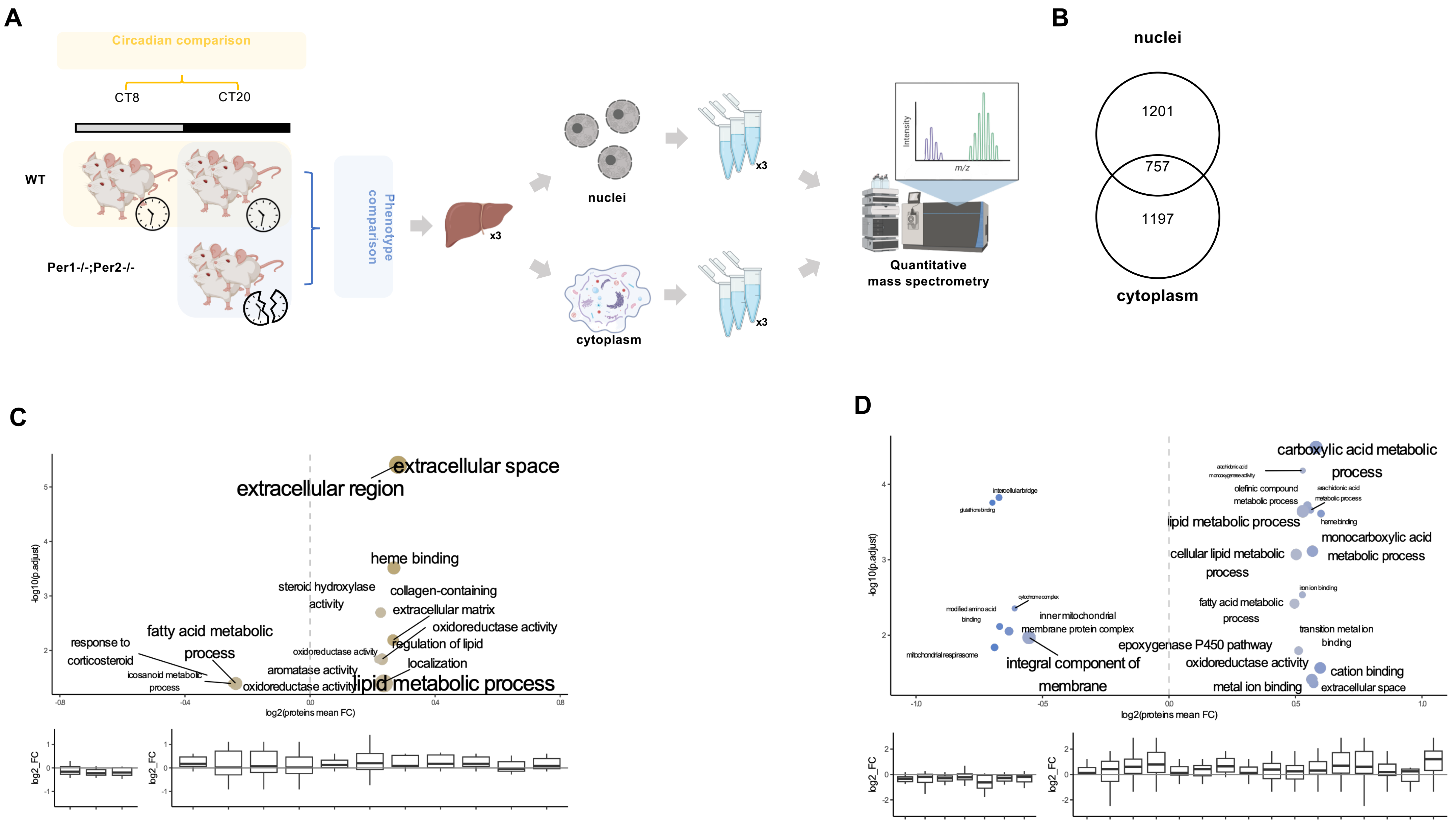


**Supp. Figure S5** *Chikhaoui, Mamgain et al*

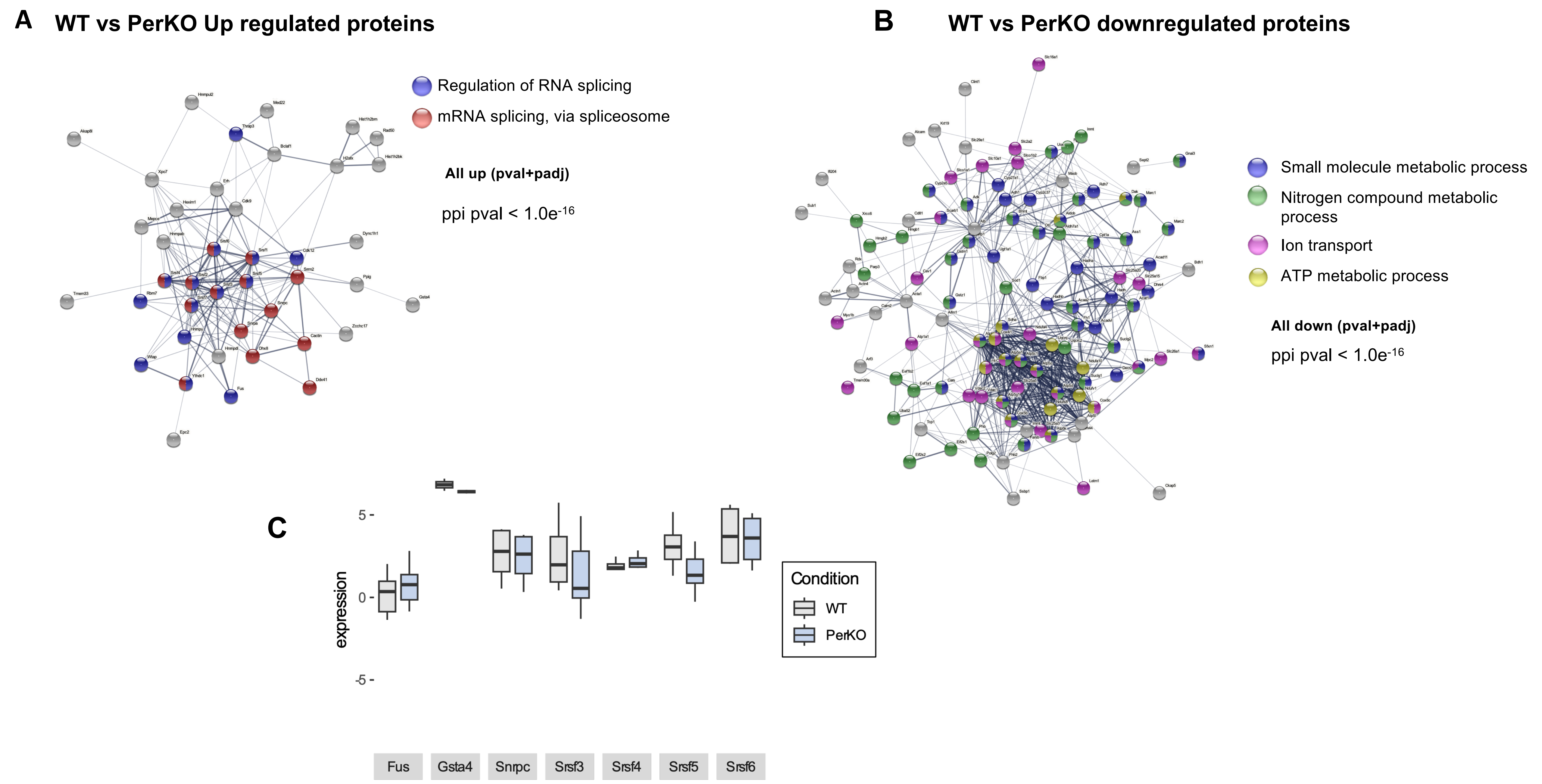


**Supp. Figure S6** *Chikhaoui, Mamgain et al*

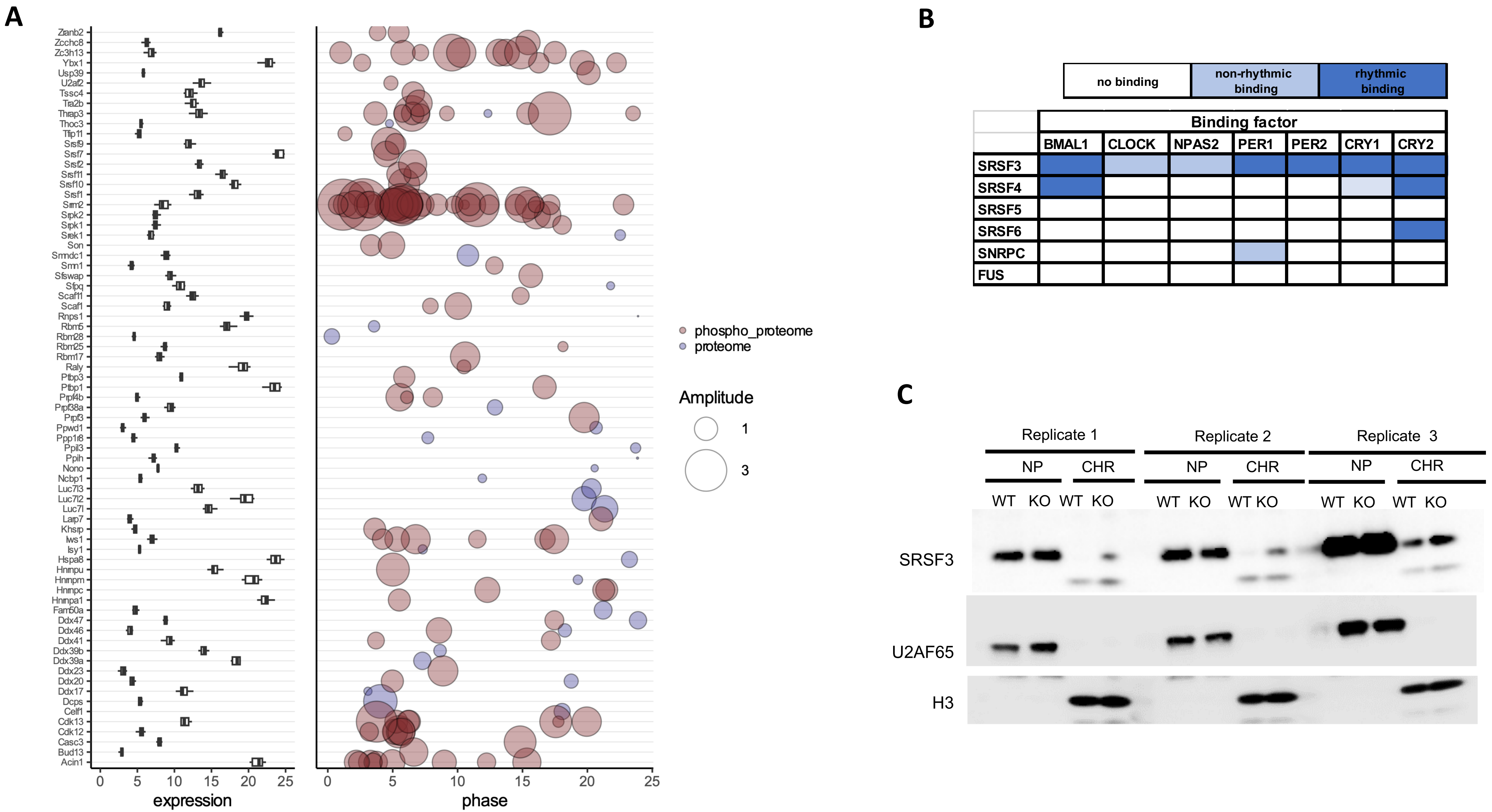


**Supp. Figure S7** *Chikhaoui, Mamgain et al*

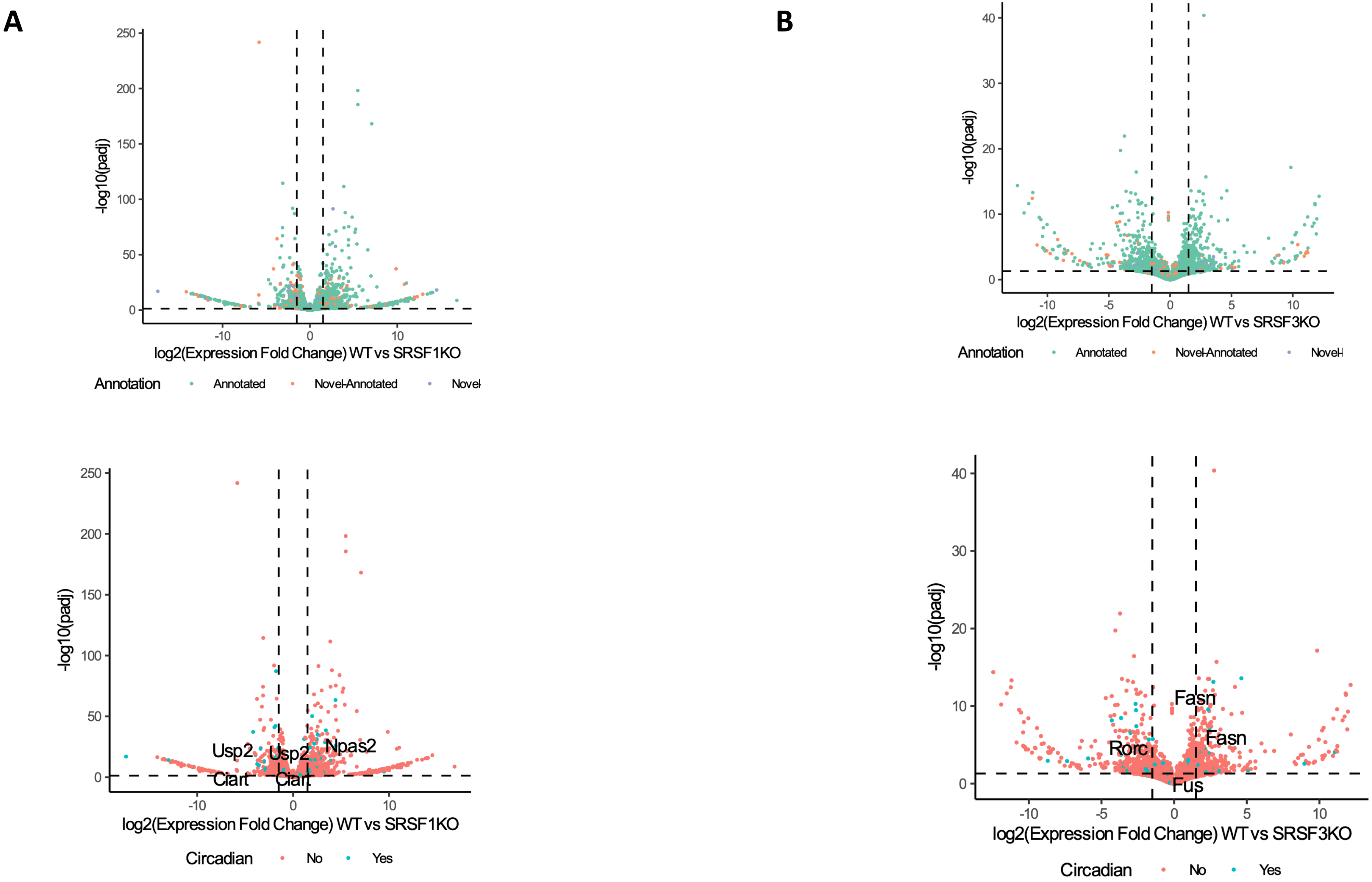


**Supp. Figure S8** *Chikhaoui, Mamgain et al*

#### Supplemental Figure Legends

##### Supplemental Figure S1

**A)** Illustration of the experimental design: briefly, WT and $Per1^{-/-}$;$Per2^{-/-}$ mice were bred in 12h light:12h dark cycles and sacrificed every 4h for WT and at CT8 or CT20 for mutants (n=2 per condition, CT = circadian time). Total RNAs were extracted from livers and sequenced using Oxford Nanopore PromethION. After base calling, sequencing data was processed to detect novel transcript, quantify isoforms and quantify splicing. Software used are in italic. **B)** Pie Chart showing fraction of total reads (all sample pooled) mapped to annotated (green), novel-annotated (orange) and novel-novel (blue) isoforms. **C)** Distribution of expression (in log2(TPM) per transcript, coloured and split by annotation class **D)** Number of transcripts that are 80% repeat, by element name. Element names separated by a comma means the element's annotations were overlapping and were merged for annotation. **E**) Expression of TE-containing transcripts, by family. **F)** Polar plot representing peak phase distribution of rhythmic productive (left) and non-productive (right) transcripts. **G)** Reactome enrichment analysis for differentially expressed transcripts (p.adjust < 0.05). Nodes are represented in yellow and genes in grey. **H)** Mosaic plot depicting distribution of rhythmic (R) and non-rhythmic (NR) in annotated/novel transcripts (left) and productive/non-productive transcripts generated from rhythmic genes (right).

##### Supplemental Figure S2

**A)** Expression patterns for individual isoforms of circadian genes exhibiting a circadian transcript usage. Shown in black are coding isoforms and in yellow non-coding transcripts. **B)** Upset plot representing the total AS landscape of the generated transcriptome when an event is included when at least one of the alternative isoform is present in the dataset. **C)** Examples of cycling AS events [event type: gene name(strand)]. Line plots showing cycling PSI values (mean for two batches) over circadian time. Boxes below show event coordinates in terms of structure (one inclusive and one alternate isoform) for each example.

##### Supplemental Figure S3

Phase plot of TE distributions (filled) at circadian AS loci compared to a randomized control set (empty). SINE (yellow), LINE (red) LTR (blue) DNA (green) TE sequences are indicated.

##### Supplemental Figure S4

**A)** Number of differential alternative splicing events in PerKO liver mice. SE: exon skipping, MX: mutually exclusive exons, RI: retained intron, A5: alternative 5' splice site, A3: alternative 3' splice site, AL: alternative last exon, AF: alternative first exon **B)** Stacked bar chart representing fraction of AS events that are unique (grey), associated with another (purple) or multiple (black) AS event of any kind on the same isoform for total and altered AS events. Fractions are shown in white.

**Supplemental Figure S5**

**A)** Experimental design. Briefly, WT and $Per1^{-/-}$;$Per2^{-/-}$ mice were bred in 12h light:12h dark cycles and sacrificed at CT8 or CT20 (n=3 per condition). Livers were collected and nuclei were separated from cytoplasm in two distinct fractions. Proteins were extracted in technical triplicates and quantitative mass spectrometry was performed to estimate protein changes between CT8 and CT20 (circadian comparison) and between WT and PerKO mice at CT20 (phenotype comparison) for each compartment **B)** Total number of proteins detected in nuclei and cytoplasm fractions and their overlap. **C)** Circadian comparison in the cytoplasm. GO Term enrichment for up and down regulated proteins in cytoplasmic fraction at CT8 compared to CT20. Bubble sizes and color correspond to number of proteins significantly mis-regulated involved in the pathway and x-axis to the mean of the identified proteins fold change. RNA expression distributions of the down and up-regulated proteins in the different pathways shown in upper panel. **D)** GO Term enrichment for up and down regulated proteins in cytoplasmic fraction at CT20 compared to PerKO. Bubble sizes and color correspond to number of proteins significantly mis-regulated involved in the pathway and x-axis to the mean of the identified proteins fold change. RNA expression distributions of the down and up-regulated proteins in the different pathways shown in upper panel.

**Supplemental Figure S6**

**A)** String network analysis on WT vs PerKO upregulated proteins **B)** String network analysis on WT vs PerKO down-regulated proteins **C)** Boxplots showing combined isoform expression for all transcripts from a given gene, list of proteins presented are differentially up-regulated in the PerKO nucleus.

**Supplemental Figure S7**

**A)** Splicing factor expression for all timepoints in WT livers are shown in the boxplot (left) and their corresponding protein (blue) or phosphorylation (red) peaks phases are shown in bubble plot (right). Size of circles corresponds to relative amplitude. Only expressed splicing factors with at least protein or one phosphorylation rhythmic are shown (Wang et al. 2017 & Robles et al. 2017). **B)**Table summarizing ChIPseq data for core-clock proteins binding to loci encoding differentially localized proteins in WT vs PerKO nuclear proteomes. Rhythmic binding of core-clock proteins (dark blue), arrhythmic binding (light blue) or no-binding (white) is indicated. **C)** Biological replicates for nucleoplasm/chromatin fractionationation, probed for SRSF3 (U2AF65 as a loading control for nucleoplasm, H3 for chromatin)

**Supplemental Figure S8**

**A)** Differential expression of Illumina RNASeq data on SRSF1 knockout liver tissue, mapped on dRNAseq transcriptome (upper panel), lower panel- differentially affected key clock genes or output genes are indicated (green dots) **B)** same as in A, for SRSF3 dataset.
